# Microbiome histidine competition mediates dietary control of systemic imidazole propionate

**DOI:** 10.64898/2026.06.22.733842

**Authors:** Paola Nol Bernardino, Christian Jacoby, Isaac T. Younker, Joshua Stemczynski, Alexander S. Little, Michael W. Mullowney, Tess H. Brunner, Joyce Ghali, Matteo Fardin, Kelsey Rose, Ramanujam Ramaswamy, Ashley M. Sidebottom, Sarah A. Tersey, Eric G. Pamer, Raghavendra G. Mirmira, Mark Mimee, Samuel H. Light

## Abstract

The gut microbiome produces numerous metabolites that influence mammalian health. While microbiome composition and diet influence metabolite concentrations, how these factors interact remains incompletely defined. Here we find production of imidazole propionate (ImP), a microbial metabolite associated with cardiometabolic and neurodegenerative diseases, is determined by the balance of competing metabolic pathways that catabolize histidine to ImP or short-chain fatty acids (SCFAs). We show glutamate serves as a preferred substrate that selectively inhibits histidine conversion to SCFAs, redirecting flux to increased ImP production across mouse- and human-derived microbial communities. We find dietary monosodium glutamate (MSG) acting via this mechanism boosts ImP production in the mouse gut, transiently impairing glucose tolerance and increasing systemic ImP. These findings show that predictable interactions between dietary substrate and microbial competition control systemic ImP levels, providing a mechanistic framework for understanding microbiome metabolite production more broadly.

## INTRODUCTION

The gut microbiome exerts broad influence over human health, with effects spanning metabolic, immune, and neurological processes.^1,2^ A key mechanism through which the microbiota interacts with its host is the production of small molecule metabolites, a subset of which are bioactive and capable of modulating host physiological processes.^1,2^ The concentrations of these metabolites vary markedly across individuals and are associated with multiple distinct health outcomes.^3,4^ Yet despite associations with diet, host genetics, and microbiome composition,^4–6^ the mechanistic basis of variation in metabolite concentrations across populations remains poorly understood.

Among microbiome-derived metabolites, imidazole propionate (ImP) has emerged as a particularly important disease-associated representative. Research by Koh *et al*.^7^ provided a foundational description of ImP, demonstrating systemic ImP levels are elevated in patients with type 2 diabetes and establishing that: (1) direct administration of ImP to mice causally impairs insulin signaling and recapitulates metabolic dysfunction, (2) these effects are mediated through activation of target of rapamycin complex 1 (mTORC1) signaling, and (3) ImP is produced from the amino acid histidine by the gut microbiota.^7^

Since this initial characterization, the association between ImP and type 2 diabetes has been independently reproduced,^8–11^ and elevated systemic ImP has been associated with a notably broad range of cardiometabolic and neuropathological conditions, including cardiovascular disease,^12–15^ Parkinson’s disease,^16^ and Alzheimer’s disease.^17^ Importantly, mouse studies using direct ImP administration established causal links between elevated systemic ImP and disease progression across these and related cardiometabolic^18–20^ and neuropathological contexts.^16,17,21^

Recent human studies demonstrate that systemic ImP levels respond to controlled dietary modification and correlate with habitual dietary patterns.^11,22^ Yet, despite these indications that diet may be involved, the factors that drive elevated ImP levels across populations remain largely unknown. Together, the broad disease relevance of ImP and this demonstrated responsiveness to diet make it an ideal model system for understanding how dietary, microbial, and host factors interact to control metabolite production in the gut microbiome.

Here, we identify competing oxidative and reductive histidine pathways in gut bacteria and demonstrate that the balance between these pathways determines ImP production in complex microbial communities. We show that dietary glutamate acts as a preferred substrate that suppresses oxidative histidine metabolism, thereby relieving competition for histidine and promoting ImP formation. In mice, dietary glutamate increases circulating ImP and impairs glucose tolerance through microbiome-dependent mechanisms, with parallel effects on systemic ImP observed in a human dietary intervention. Together, these findings demonstrate that microbial substrate preference hierarchies create predictable links between dietary composition and metabolite output, providing a mechanistic framework for diet-microbiome-metabolite interactions.

## RESULTS

### A Stickland-like reductive histidine pathway converts histidine to ImP

Previous studies have shown that the gut microbiome produces ImP from the amino acid histidine.^7^ Bacteria encoding urocanate reductase (UrdA) convert urocanate, a histidine degradation product, to ImP^7,16,21,23,24^ but, to our knowledge, gut bacteria capable of fully transforming histidine to ImP have not been previously characterized (**Figure 1A**). We hypothesized that species required both *urdA* and histidine ammonia-lyase (*hutH*), which deaminates histidine to urocanate, to convert histidine to ImP (**Figure 1A**). To test this, we selected six gut bacterial species that encoded either *hutH* alone (+*hutH*/-*urdA*), *urdA* alone (-*hutH*/+*urdA*), or both genes (+*hutH*/+*urdA*). Consistent with our hypothesis, we found all *urdA*-encoding strains converted urocanate to ImP, but only *Fusobacterium_A ulcerans* and *Parasutterella excrementihominis* strains encoding both *hutH* and *urdA* converted histidine-^13^C_6_ to ^13^C_6_-ImP (**Figure 1B**).

**Figure 1.**
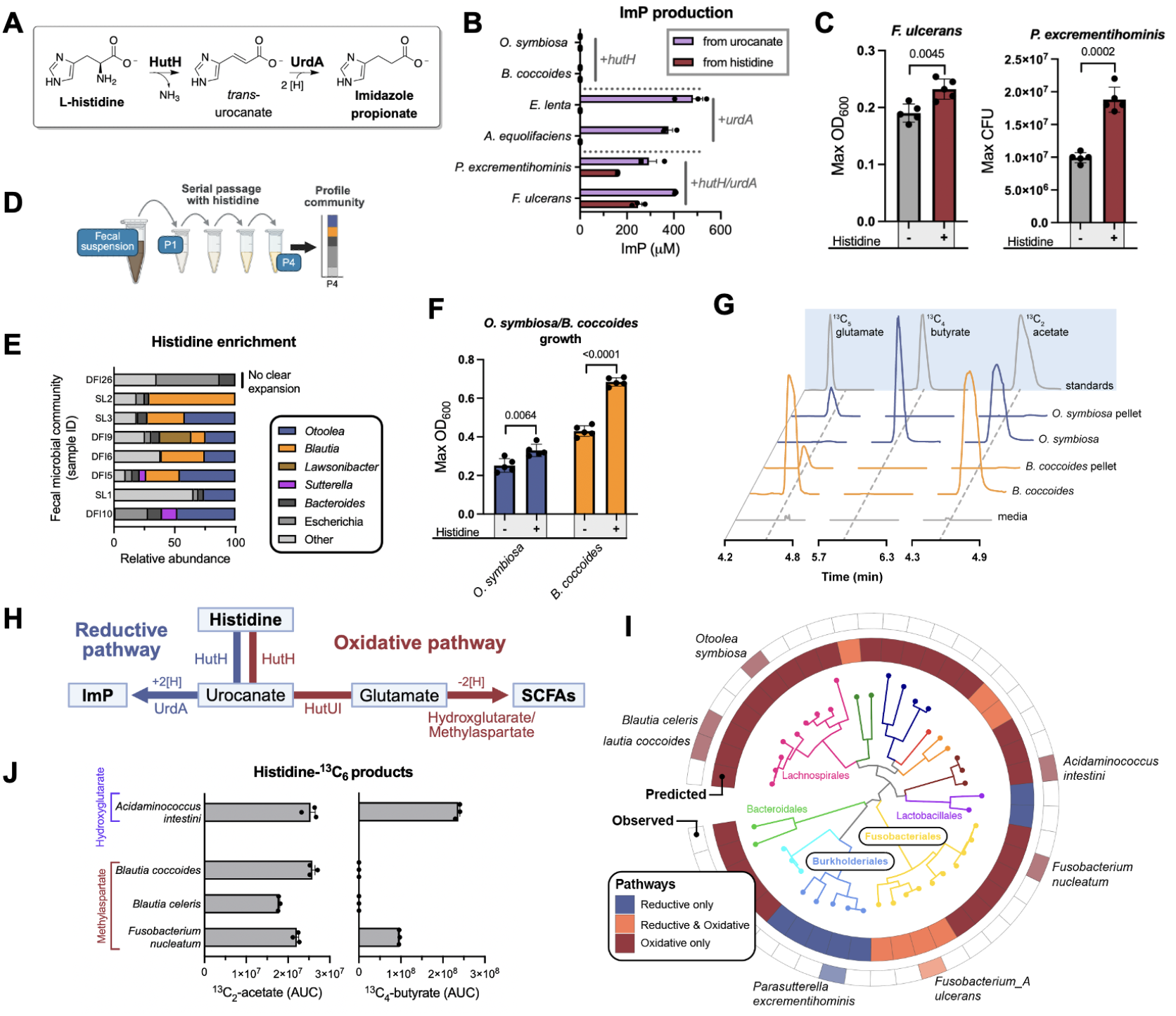
Gut bacteria catabolize histidine via distinct Stickland-like oxidative and reductive pathways. **(A)** Enzymatic steps in histidine conversion to ImP. **(B)** ImP production from urocanate or histidine as determined by LC-MS measurement of ImP isotopologues in urocanate-^12^C_6_ and histidine-^13^C_6_ supplemented culture media. (**C**) Maximum cell density of *F. ulcerans* and colony forming units (CFU) of *P. excrementihominis* cultured with or without 10 mM histidine supplementation. (**D**) Experimental design for enriching histidine-catabolizing bacteria from human fecal communities through serial passaging with urocanate as the primary substrate. (**E**) Community composition of human fecal microbial communities following histidine enrichment passaging. (**F**) Maximum cell density of *O. symbiosa* and *B. coccoides* strains grown with or without 10 mM histidine supplementation. (**G**) Representative LC-MS chromatogram showing derivatized metabolites detected in indicated cell cultures supplemented with histidine-^13^C_6_. (**H**) Overview of Stickland-like reductive and oxidative histidine pathways that convert histidine to ImP or SCFAs. (**I**) Phylogenetic distribution of predicted reductive and oxidative histidine pathways across representative gut bacterial genomes. Outer rings indicate pathway presence based on marker gene identification of species with experimentally confirmed pathways. (**J**) Production of short-chain fatty acids (SCFAs) from histidine-^13^C_6_ by oxidative pathway-containing bacteria, as determined by ^13^C-metabolite tracing. Data shown as mean ± SEM. Statistical analyses by Two-tailed Welch’s t-test.

To test whether the identified histidine-metabolizing strains produce ImP in the gut, we colonized germ-free mice with *F. ulcerans, P. excrementihominis*, or *Escherichia coli* (**Figure S1A**). We found that both histidine-metabolizing strains resulted in elevated cecal and blood ImP, whereas the non-histidine-metabolizing control strain *E. coli* resembled the germ-free control (**Figure S1B & S1C**). Similarly, colonization with *F. ulcerans* or *P. excrementihominis* impaired glucose tolerance, comparable to the previously reported impact of direct ImP administration (**Figure S1D**),^7^ while *E. coli* colonization had no effect. These findings demonstrate that *F. ulcerans* and *P. excrementihominis* convert histidine to ImP in the gut and reproduce metabolic dysfunction associated with elevated ImP.

We next sought to clarify the function of histidine conversion to ImP in *F. ulcerans* and *P. excrementihominis*. Notably, the HutH-UrdA reactions bear strong resemblance to characterized reductive Stickland fermentation pathways that metabolize aromatic amino acids through α,β-unsaturated acrylate intermediates to derivative propionate products (**Figure S2**).^25^ As *F. ulcerans* and *P. excrementihominis* are obligate anaerobes and reductive Stickland pathways enable a subset of anaerobic bacteria to fuel growth on amino acids,^26,27^ we reasoned the HutH-UrdA pathway might possess an analogous function. Consistent with this hypothesis, we found supplementing media with 10 mM histidine promoted *F. ulcerans* and *P. excrementihominis* growth (**Figure 1C**). Together, these structural and functional similarities suggest that the HutH-UrdA pathway represents a “Stickland-like” reductive histidine metabolism present in a subset of gut bacteria.

### Stickland-like oxidative histidine pathways metabolize histidine via glutamate to SCFAs

Anaerobic bacteria classically employ Stickland fermentation to catabolize pairs of amino acids and other substrates through a net redox neutral process that couples the oxidation of one substrate with the reduction of another.^26,27^ Different bacterial species catabolize the same amino acid substrate through discrete reductive or oxidative Stickland pathways that generate distinct terminal products.^25,28^ Based on this precedent, we asked whether a Stickland-like oxidative histidine pathway might also operate in the gut.

To identify bacteria with a Stickland-like oxidative histidine pathway, we applied histidine-supplemented selective growth media to enrich bacteria from eight human fecal microbial communities (**Figure 1D**). We sequenced resulting enrichment communities and found that members of two Clostridial genera, *Otoolea* and *Blautia*, consistently expanded in the presence of histidine (**Figure 1E**). We selected *Otoolea symbiosa* MSK.7.21 and *Blautia coccoides* UOC.14.97 as representative of these genera and confirmed supplementation with 10 mM histidine promoted growth of both strains (**Figure 1F**).

To address the basis of histidine metabolism in *O. symbiosa* and *B. coccoides*, we incubated each strain with histidine-^13^C_6_ and found they efficiently consumed histidine-^13^C_6_ without producing ImP (**Figure 1B & 1G**). Instead, both strains converted histidine to short-chain fatty acids (SCFAs), with *B. coccoides* producing ^13^C_2_-acetate and *O. symbiosa* producing both ^13^C_2_-acetate and ^13^C_4_-butyrate (**Figure 1G**).

To gain insight into the mechanism of histidine catabolism to SCFAs, we conducted transcriptomics analysis of *O. symbiosa* cells with histidine as the primary growth substrate. We observed Hut pathway genes *hutH, hutU*, and *hutI*, which catalyze the conversion of histidine to glutamate,^29^ were among the most highly expressed in this condition (**Figure 1H, Figure S3A-S3B, Table S1**). As *O. symbiosa* was previously reported to ferment glutamate via the oxidative 2-hydroxyglutarate pathway and *B. coccoides* encodes the methylaspartate pathway, a second Stickland-like pathway responsible for oxidative glutamate fermentation in bacteria,^30–32^ we reasoned the observed Hut pathway expression could be indicative of oxidative histidine catabolism via a glutamate intermediate.

To test this pathway model, we examined utilization of pathway intermediates. Consistent with the Hut pathway mediating histidine catabolism, supplementation with 10 mM urocanate (the first pathway intermediate^29^) promoted growth of both *O. symbiosa* and *B. coccoides* and was efficiently consumed, comparable to histidine (**Figure S3C-S3F**). To confirm glutamate as the downstream intermediate, we performed histidine-^13^C_6_ tracing and observed both species produced ^13^C_5_-glutamate (**Figure 1G**), consistent with conversion through the Hut pathway. Together with the transcriptional evidence, these results provide evidence that *O. symbiosa* and *B. coccoides* catabolize histidine through a Stickland-like oxidative pathway that channels substrate into oxidative glutamate fermentation pathways.

### Reductive and oxidative histidine pathways are widespread and genetically co-integrated in a subset of gut bacteria

Having identified distinct Stickland-like reductive and oxidative catabolic pathways that convert histidine to ImP or SCFAs (**Figure 1H**), we sought to clarify the distribution of these pathways across gut bacteria. We performed comparative genomics analysis on a collection of 4,744 representative human gut prokaryote genomes and metagenome-assembled genomes to identify species with potential oxidative and reductive histidine pathways.^33^

These analyses identified predicted oxidative histidine pathways in 169 gut bacteria species spanning 51 genera. (**Figure 1I, Table S2**). Consistent with this broad predicted taxonomic distribution, we confirmed that additional strains encoding the oxidative histidine pathway from three distinct phyla efficiently converted histidine-^13^C_6_ to ^13^C-labeled SCFAs (**Figure 1J**). Further supporting the conclusion that these predictions capture the dominant histidine-catabolizing pathway in gut bacteria, we found predicted oxidative histidine pathways were strongly associated with histidine consumption in culture media metabolomes previously reported for 135 gut bacteria isolates (**Figure S4, Table S3**).^34^ These results thus provide evidence the oxidative histidine pathway represents a key histidine catabolic pathway within taxonomically distinct gut bacteria.

Comparative genomics analysis of predicted reductive histidine metabolism identified the pathway in 31 species and 17 genera, the majority of which resolved to Fusobacteriaceae or Burkholderiaceae clades that included the *F. ulcerans* and *P. excrementihominis* strains with experimentally validated activity, respectively (**Figure 1I, Table S2**). Notably, scrutiny of these reductive histidine pathway clades showed the oxidative histidine pathway was conserved in Fusobacteriaceae but absent from Burkholderiaceae genomes (**Figure S5A**). This distinct relationship between the two histidine pathways was further reflected in the genetic organization of reductive histidine pathway genes across these two clades, with *hutH* co-localizing with *urdA* in Burkholderiaceae genomes but oxidative histidine pathway genes *hutUI* in Fusobacteriaceae genomes (**Figure S5B**). Consistent with these observations, we confirmed *P. excrementihominis* exclusively converted histidine-^13^C_6_ into ^13^C_6_-ImP, whereas *F. ulcerans* generated both reductive and oxidative histidine pathway products (**Figure S5C**). These studies thus reveal that reductive and oxidative histidine pathways are widespread and exhibit distinct patterns of genetic co-integration across gut bacteria, with potential functional ramifications for histidine metabolism.

### The oxidative histidine pathway dominates direct histidine competition, reduces ImP production in microbial communities, and improves glucose tolerance in mice

Having identified oxidative and reductive histidine pathways, we next sought to clarify their representation within the gut microbiome. To address this, we analyzed the abundance of bacteria with oxidative and reductive histidine pathways across existing human gut metagenomes (**Table S4**). We found oxidative pathway bacteria substantially outnumbered reductive pathway bacteria in both species richness (i.e., number of species encoding the pathway) and cumulative relative abundance across samples (**Figure S6A & S6B**).

Considering the high relative abundance of bacteria with the oxidative histidine pathway across human microbiomes, we wondered whether competition from the oxidative histidine pathway activity could suppress ImP production. To address this hypothesis, we cultured *F. ulcerans* and *P. excrementihominis* with or without competing species possessing the oxidative histidine pathway. Indeed, we found addition of bacteria with the oxidative histidine pathway led to a marked decrease in ^13^C_6_-ImP produced from histidine-^13^C_6_ (**Figure 2A**). These results demonstrate that oxidative histidine pathway bacteria establish metabolic competition for histidine that suppresses ImP production by reductive pathway species.

**Figure 2.**
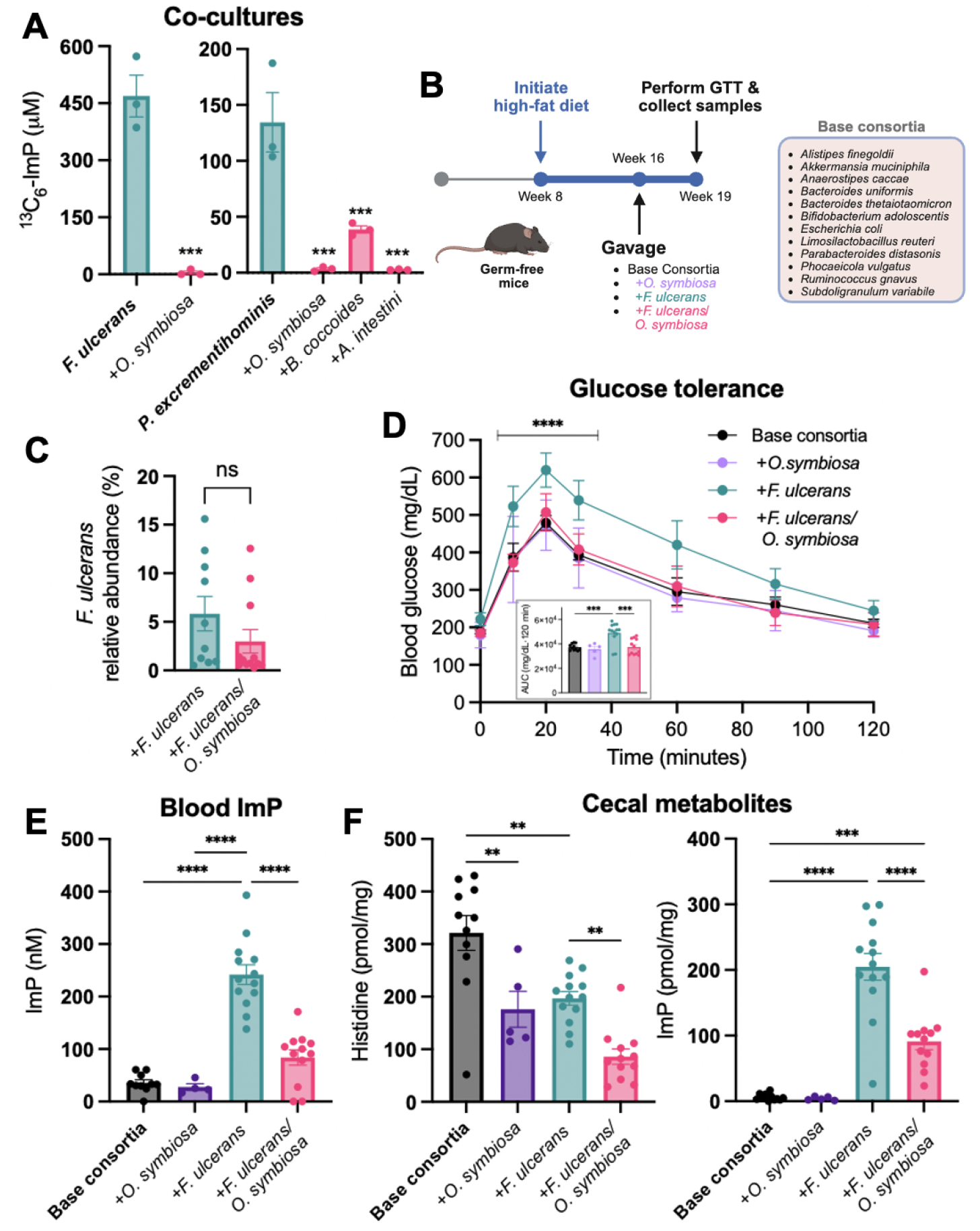
The oxidative histidine pathway dominates direct histidine competition and limits systemic ImP levels in mice. **(A)** ^13^C_6_-ImP production from reductive pathway bacteria cultured alone or with oxidative pathway bacteria in histidine-^13^C_6_ supplemented media. **(B)** Experimental design for HFD-consortia gnotobiotic mouse model. **(C)** Fecal *F. ulcerans* relative abundance within HFD consortia determined by 16S rRNA sequencing. **(D)** Glucose tolerance, **(E)** blood ImP and **(F)** cecal metabolite concentrations in consortia colonized mice. Data shown as mean ± SEM. Statistical analyses by Two-tailed Welch’s t-test or one-way ANOVA with Tukey’s post-hoc test.

To determine whether the observed effect of histidine competition in suppressing ImP production translated to an *in vivo* context relevant to cardiometabolic disease, we next developed an “HFD-consortia mouse model,” which involved placing germ-free mice on a high-fat diet for 8 weeks, followed by colonization with a previously characterized defined “base consortia” comprising 12 human gut bacterial species selected to capture much of the taxonomic and metabolic diversity of a complex microbiome while lacking histidine metabolic pathways (**Figure 2B**).^35^

After confirming addition of *F. ulcerans* to the base consortia recapitulated previously observed blood ImP and glucose tolerance phenotypes, we tested the impact of histidine competition by introducing *O. symbiosa* to this model. We found that *O. symbiosa* counteracted the effect of *F. ulcerans* on blood ImP and glucose response, without substantially impacting *F. ulcerans* abundance (**Figure 2C-2E**). Consistent with histidine competition playing a key role in these phenotypes, *O. symbiosa* colonization had little effect on ImP and glucose response in the absence of *F. ulcerans* but resulted in lower cecal histidine and ImP concentrations in its presence (**Figure 2F**). These results thus demonstrate metabolic competition for histidine within the gut can reduce systemic ImP and associated cardiometabolic phenotypes.

### Glutamate is a preferred growth substrate that inhibits the oxidative histidine pathway in diverse gut bacteria

As our *in vitro* and gnotobiotic studies established that the oxidative histidine pathway imposes metabolic competition that suppresses ImP production in microbial communities, we reasoned low oxidative histidine pathway activity might contribute to the elevated ImP levels observed in cardiometabolic disease patients. We first considered whether this might arise from a scenario similar to our HFD-consortia gnotobiotic studies, in which bacteria with the oxidative histidine pathway were underrepresented. However, analysis of gut metagenomes from cardiometabolic disease patients revealed that bacteria encoding the oxidative histidine pathway were nearly universally present and exhibited comparable relative abundances to healthy individuals, arguing against underrepresentation of bacteria with the oxidative histidine pathway as a meaningful contributor to elevated ImP in disease (**Figure S7A & S7B, Table S4)**.

We next considered whether low oxidative histidine pathway levels might result from factors that inhibit oxidative activity irrespective of pathway abundance. Our earlier characterization of the oxidative histidine pathway showed it channels histidine into oxidative glutamate fermentation pathways **(Figure 1H)**, with the presumed energy-yielding portion occurring downstream of a glutamate intermediate. Consequently, direct glutamate entry into the pathway could bypass the initial non-energy-yielding steps of histidine conversion, possibly serving as a more efficient substrate to bacteria with the oxidative histidine pathway **(Figure 3A)**. As bacteria tend to preferentially metabolize substrates that net higher energy yields,^36^ we reasoned dietary glutamate might thus provide a preferred growth substrate capable of suppressing the oxidative histidine pathway.

**Figure 3.**
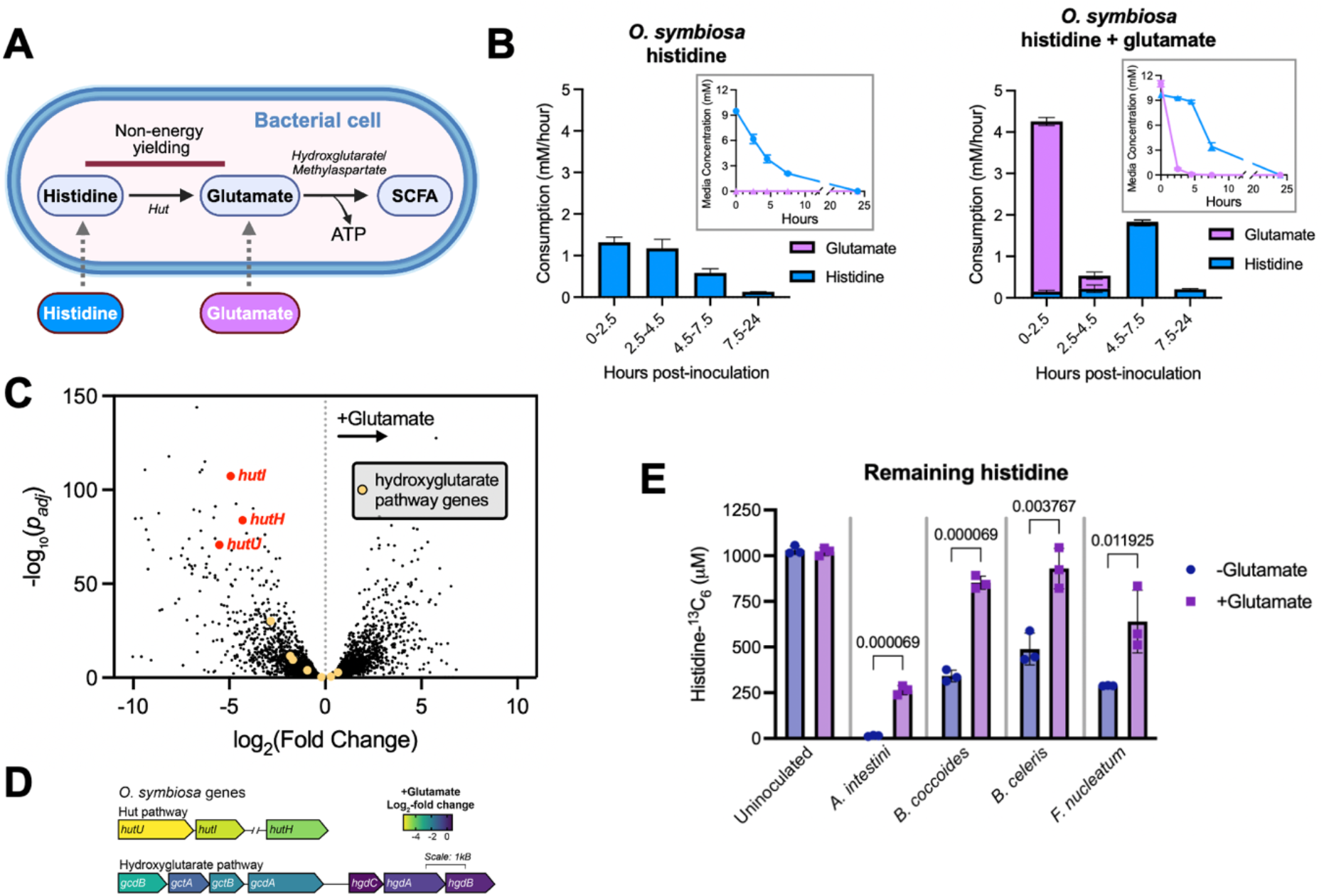
Glutamate is a preferred substrate that suppresses oxidative histidine pathway activity in gut bacteria. **(A)** Bacterial cell illustration highlighting how downstream glutamate entry into the oxidative pathway could make it an efficient substrate. **(B)** Time course of glutamate and histidine consumption rates in *O. symbiosa* cultures supplemented with 10 mM histidine ± 10 mM glutamate. Insets show culture histidine and glutamate concentrations. **(C)** Differential gene expression in *O. symbiosa* with histidine ± glutamate. **(D)** Fold-change of *O. symbiosa* oxidative histidine pathway gene expression in the presence of glutamate. **(E)** Production of ^13^C-labeled oxidative pathway products (acetate and butyrate) from cultures supplemented with histidine-^13^C_6_ ± glutamate. Data shown as mean ± SEM. Statistical analyses by Two-tailed Welch’s t-test.

To address the impact of glutamate on oxidative histidine metabolism, we tracked *O. symbiosa* growth and amino acid consumption in cultures with or without 8 mM glutamate supplementation. We found supplemented glutamate was rapidly depleted from the media over the first 4.5 hours **(Figure 3B)**. In this interval, glutamate availability was associated with a marked suppression of *O. symbiosa* histidine-^13^C_6_ utilization **(Figure 3B)**. However, following the near complete consumption of glutamate by 4.5 hours, the rate of histidine-^13^C_6_ consumption recovered **(Figure 3B)**. These results thus provide evidence that *O. symbiosa* prioritizes glutamate metabolism, suppressing histidine utilization until this preferred substrate is depleted.

To clarify the basis of glutamate-mediated suppression of histidine metabolism, we performed transcriptomic analysis of *O. symbiosa* cells cultured with or without glutamate supplementation. We found glutamate inhibited the transcription of the *hutHUI* genes responsible for converting histidine to glutamate, but not the 2-hydroxyglutarate pathway genes required for glutamate oxidation **(Figure 3C & 3D, Table S5)**. These findings demonstrate that glutamate transcriptionally suppresses histidine catabolism in a manner consistent with characterized hierarchical substrate utilization mechanisms that prioritize energetically favorable substrates in bacteria.^36^

Reasoning that the logic of glutamate’s energetic favorability making it a preferred substrate extended to other bacteria with the oxidative histidine pathway, we expanded our analysis of the effect of glutamate to four additional species (*Acidaminococcus intestini, B. coccoides, B. celeris, and Fusobacterium nucleatum*) that span the taxonomic diversity of the oxidative histidine pathway in gut bacteria. We found supplemented glutamate was efficiently consumed and that this was associated with decreased histidine consumption across the tested species **(Figure 3E, Figure S8)**. These results demonstrate that glutamate substrate preference is conserved across phylogenetically diverse bacteria encoding the oxidative histidine pathway.

### Glutamate decreases histidine competition to promote ImP production across microbial communities

We next sought to address the effect of glutamate-mediated oxidative histidine pathway suppression on histidine competition within microbial communities. To determine whether glutamate inhibition could counteract the observed effect of *between-species* histidine competition in limiting ImP production (**Figure 4A**), we tested *P. excrementihominis* histidine-^13^C_6_ metabolism in co-culture with the oxidative histidine pathway bacteria *O. symbiosa, B. coccoides*, or *A. intestini*. Consistent with the hypothesis, we found the addition of glutamate partially or fully restored ^13^C_6_-ImP production in the presence of these strains (**Figure 4B**).

**Figure 4.**
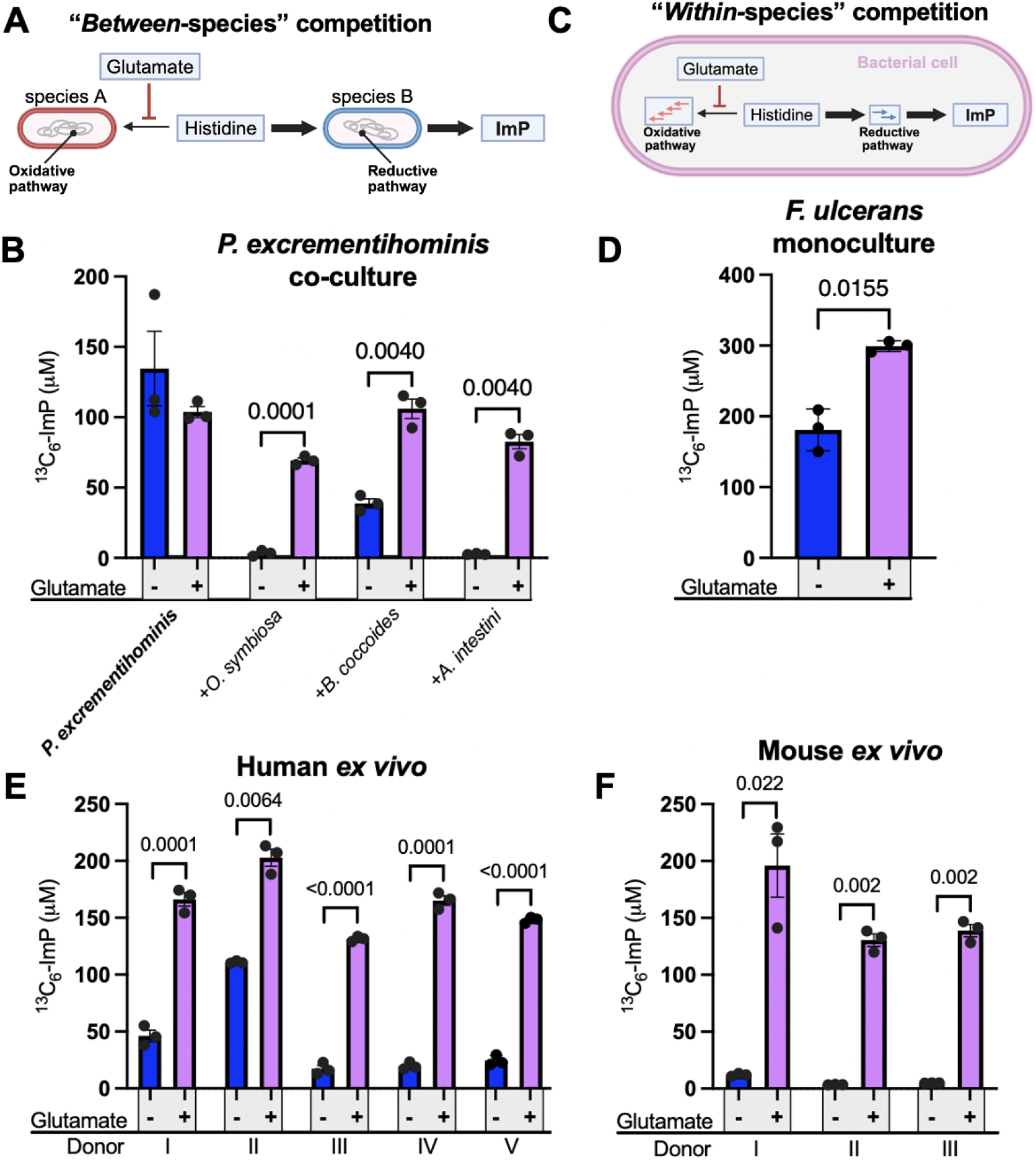
Glutamate-mediated suppression of the oxidative histidine pathway promotes ImP production across microbial communities. **(A)** Model of proposed impact of glutamate on *between*-species histidine competition. **(B)** ^13^C_6_-ImP production from *P. excrementihominis* cultures supplemented with histidine-^13^C_6_ ± glutamate in the presence or absence of oxidative histidine pathway bacteria. **(C)** Model of proposed impact of glutamate on *within*-species histidine competition. **(D**) ^13^C_6_-ImP production from *F. ulcerans* cultures supplemented with histidine-^13^C_6_ ± glutamate. **(E)** ^13^C_6_-ImP produced from *ex vivo* human fecal suspensions supplemented with histidine-^13^C_6_ ± glutamate. **(F)** ^13^C_6_-ImP produced from *ex vivo* mouse fecal suspensions supplemented with histidine-^13^C_6_ ± glutamate. Data shown as mean ± SEM. Statistical analyses by Two-tailed Welch’s t-test.

Beyond *between-species* competition, we reasoned that suppressed oxidative histidine metabolism might also relieve histidine competition within individual cells encoding both oxidative and reductive histidine pathways, thereby promoting ImP production through a distinct *within-species* mechanism (**Figure 4C**). To address the effect of glutamate on *within-species* histidine competition, we examined glutamate’s impact on histidine-^13^C_6_ metabolism in *F. ulcerans*, which encodes both oxidative and reductive histidine pathways. Consistent with glutamate also affecting *within*-species histidine competition, we found glutamate addition nearly doubled ImP production in *F. ulcerans* cultures (**Figure 4D**).

Having established the relevance of glutamate for oxidative pathway suppression in multiple organisms and both *between-* and *within*-species histidine competition, we reasoned glutamate might promote ImP production across compositionally diverse natural gut microbial communities. Toaddress this hypothesis, we tested the effect of glutamate on *ex vivo* histidine-^13^C_6_ metabolism in fecal suspensions collected from 5 human and 3 mouse donors. We found glutamate was efficiently consumed and resulted in elevated histidine and ImP levels, consistent with increased reductive histidine metabolism coming at the expense of oxidative pathway activity (**Figure 4E & 4F, Figure S9A-S9D**). These studies thus demonstrate that glutamate promotes ImP production in complex fecal communities through a mechanism that transcends interindividual and interspecies variation in microbiome composition.

### Dietary MSG modulates microbial histidine competition, increases circulating ImP, and impairs glucose tolerance

Intestinal glutamate originates from multiple dietary sources, including digestive protein hydrolysis and monosodium glutamate (MSG), a common flavor enhancer consumed at variable rates across cuisines.^37,38^ Notably, a previous study demonstrated that dietary MSG induces cardiometabolic dysfunction, including impaired glucose tolerance, in mice.^39^

Because (1) MSG provides a defined source of glutamate independent of the complex amino acid mixtures generated during protein digestion and (2) elevated ImP could plausibly explain previously reported MSG-associated cardiometabolic phenotypes, we selected dietary MSG supplemented with histidine to test whether glutamate availability modulates microbial histidine pathway competition and increases systemic ImP in vivo.

As the conserved effect of glutamate on ImP production in mouse fecal suspensions suggested specific pathogen-free (SPF) mice would provide a biologically relevant system for testing dietary MSG effects, we assessed the impact of MSG supplementation through drinking water in SPF mice maintained on a high-fat diet (**Figure 5A**). We found that dietary MSG resulted in elevated blood ImP and impaired glucose tolerance compared to a sodium-matched (NaCl) control (**Figure 5B & 5C**). Consistent with these phenotypes resulting from the effect of dietary glutamate on histidine pathway competition within the distal gastrointestinal tract rather than broad physiological or microbial restructuring, MSG treatment led to elevated cecal ImP (**Figure 5D**) with no major effects on mouse body weight, water consumption, or overall gut microbiome composition (**Figure S10A-S10D**).

**Figure 5.**
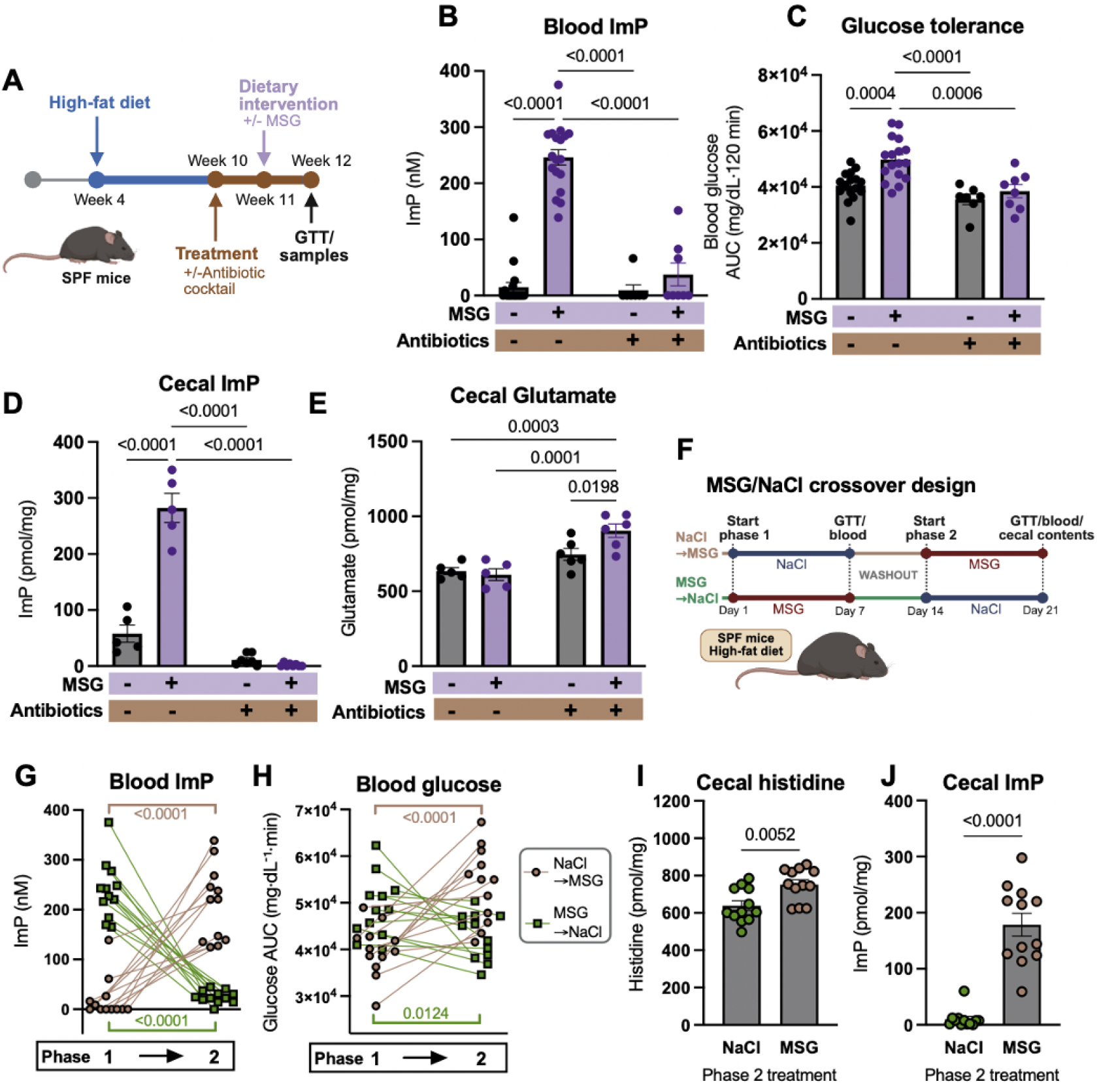
Dietary MSG transiently increases circulating ImP and impairs glucose tolerance in SPF mice through microbiome-dependent mechanisms. **(A)** Experimental design for mouse dietary MSG study. **(B)** Blood ImP, **(C)** glucose tolerance, **(D)** cecal ImP, and **(E)** cecal glutamate from dietary MSG study, analyzed by one-way ANOVA. **(F)** Experimental design for mouse MSG crossover study. **(G)** Blood ImP and **(H)** glucose tolerance from individual mice in crossover study, analyzed by paired t-test. **(I)** Cecal histidine and **(J)** cecal ImP following phase 2 of mouse crossover study, analyzed by Two-tailed Welch’s t-test.

To further corroborate the role of the microbiome in observed phenotypes, we tested the effect of a microbiome-inhibiting antibiotic cocktail on the high MSG diet. Consistent with dietary glutamate serving as a microbial substrate in the distal gastrointestinal tract, we found that combined MSG and antibiotic treatment led to elevated cecal glutamate levels (**Figure 5E**). We further found antibiotic treatment depleted the gut microbiome and abrogated the effect of MSG on cecal ImP, blood ImP, and glucose tolerance (**Figure 5B-5D, Figure S10**). These findings demonstrate that dietary MSG elevates ImP and impairs glucose tolerance through microbiome-dependent mechanisms, consistent with direct glutamate suppression of the oxidative histidine pathway activity in the distal gut.

Because the mechanism of glutamate-induced ImP production suggested by our studies involves a transcriptionally mediated change in histidine competition rather than a stable shift in microbiome composition, we next asked whether MSG phenotypes could be reversed by dietary intervention. We conducted a crossover experiment in which mice received MSG or sodium-matched diets before swapping diets (**Figure 5F**). We found blood ImP dropped and glucose tolerance improved following removal of MSG from the diet (**Figure 5G & 5H**). Consistent with these changes reflecting increased histidine competition in the gut, MSG cessation was associated with reduced cecal histidine and ImP (**Figure 5I & 5J**). These results demonstrate an acute requirement for dietary glutamate to maintain elevated ImP and establish that associated metabolic phenotypes are reversible through dietary modification.

## DISCUSSION

The studies presented here demonstrate that ImP production in gut microbial communities is governed by competition between oxidative and reductive histidine pathways, and that this competition is predictably modulated by dietary glutamate acting through conserved substrate preference hierarchies. In mice, dietary MSG increases circulating ImP and impairs glucose tolerance through microbiome-dependent mechanisms, with parallel effects on systemic ImP observed in a human dietary intervention. These findings establish a mechanistic link between a specific dietary input, microbial pathway competition, and a disease-relevant circulating metabolite.

A central insight of this work concerns the mechanistic determinants of metabolite levels. Traditional approaches have sought to predict metabolite abundance from taxonomic or genomic features, yet these correlations show limited consistency across populations.^4,6^ Our findings suggest that incorporating pathway competition may better explain variation in microbial metabolite production across the population.

Our work further demonstrates that the balance between competing pathways can be predictably modulated by dietary factors acting on substrate preference hierarchies. While the generalizability of this principle to other metabolites remains to be established, substrate preference hierarchies are often conserved across bacterial taxa,^36^ and dietary fiber was previously shown to suppress circulating indoxyl sulfate levels,^40^ presumably by providing a preferred substrate that diverts tryptophan metabolism away from indole production. Together, these observations suggest that accounting for dietary interactions with conserved substrate preference hierarchies may provide a general framework for predicting diet-microbiome-metabolite interactions.

In addition, these findings raise the possibility that dietary modulation of ImP production could offer therapeutic benefit in cardiometabolic and neuropathological conditions in which ImP has been implicated as a causal contributor. Several considerations limit direct translation, however. A causal role for ImP in cardiometabolic dysfunction has been established in mice, but whether comparable effects occur in humans remains unknown. Furthermore, MSG represents a defined and concentrated glutamate source that differs meaningfully from the complex amino acid mixtures generated during protein digestion, leaving open the question of which dietary components most meaningfully drive elevated ImP in at-risk individuals. As histidine availability, microbiome composition, and other dietary factors likely interact with glutamate to shape ImP production in ways the present studies do not fully resolve, dedicated human intervention studies will be required to identify dietary components that can reduce ImP levels to meaningful metabolic benefit.

Ultimately, this work advances a mechanistic framework for understanding how diet and microbial pathway competition jointly determine the levels of disease-relevant metabolites. This framework may improve predictions of microbiome output across individuals and dietary contexts and inform the development of precision nutrition strategies aimed at selectively modulating specific microbial metabolites for therapeutic effect.

## METHODS

### Bacterial strains and culture media

Bacterial strains used in this study were acquired the Duchossois Family Institute (DFI) Biobank or other reputable strain collections.^41,42^ Unless otherwise specified, all references to bacterial species throughout the manuscript refer to the specific strain listed in **Table S7**. Strains were routinely cultured under anaerobic conditions in a Coy vinyl anaerobic chamber (2-5% H_2_, 2-5% CO_2_, N_2_ balance) at 37°C. Growth media included Brain Heart Infusion Medium with Hemin (BHIM), basal, and chemically defined media. The chemically defined medium contained 1 g/L NH_4_Cl, 6 g/L Na_2_HPO_4_, 3 g/L KH_2_PO_4_, 0.5 g/L NaCl, 14.7 mg/L CaCl_2_, 246 mg/L MgSO_4_·7H_2_O, 0.5% glucose, 0.05% L-cysteine, 2.5 mg/L vitamin K3, 2 mg/L FeSO_4_, 5 mg/L hemin, 5 ng/mL vitamin B12.

### Genomic and bioinformatic analyses

#### Identification of histidine pathway genes in gut bacterial genomes

To identify bacterial species encoding candidate histidine metabolic pathways, characterized enzymes catalyzing key pathway steps (**Table S6**) were used as BLASTp^43^ (v2.9.0) queries against a custom database of protein sequences from prevalent UHGG^33^ (v2.0.1) species representatives. Significant hits were identified using an e-value cutoff < 1E-5 with enzyme-specific percent identity and coverage thresholds applied to filter results (**Table S6**). Species were classified as encoding the reductive pathway if homologs for both *hutH* and *urdA* were identified. Species were classified as encoding the oxidative pathway if homologs for the hut (*hutH* and *hutU*) and methylaspartate (*mal*) or hydroxyglutarate (*gcdA*) pathways were identified. Results from these analyses are provided in **Table S2**.

#### Phylogenetic analysis of histidine-metabolizing taxa

Single copy core genes for all UHGG (v.2.01) species representatives were identified using hmmsearch^44^ (v3.1b2; evalue cutoff 1e^-5) with the Bac120 set of profile hidden Markov models from GTDB-Tk^45^ (v2-r220). A subset of taxa with the predicted oxidative and reductive histidine metabolism pathways was selected and the 27 most prevalent single copy marker proteins (present in at least 42/43 genomes) were aligned with mafft (v7.505; --retree 2 – maxiterate 0).^46^ Aligned sequences were trimmed with TrimAI (v1.4.1; -gappyout),^47^ concatenated, and a phylogenetic tree inferred with FastTree (v2.1.11; default settings).^48^ The phylogenetic tree was midpoint rooted and visualized using the phytools^49^ (v2.4.4) and ggtree^50^ (v3.6.2) suite of packages in R (v4.2.2).^51^

### *In vitro* bacterial assays

#### Bacterial histidine and urocanate growth

*O. symbiosa, B. coccoides*, and *F. ulcerans* growth at 37°C was monitored by optical density at 600 nm (OD_600_) every 20 minutes for 48 hours using a LogPhase 600 plate reader (BioTek) with LogPhase 600 App software (v1.07) with continuous shaking (200 rpm) under anaerobic conditions in base media ± 10 mM L-histidine or urocanate.

#### P. excrementihominis growth was assessed by CFU enumeration

Cells were grown in Gifu Anaerobic Medium (GAM) at 37°C, harvested by centrifugation, washed in anaerobic PBS, and resuspended to normalize inoculum. Cultures were inoculated into GAM alone, GAM + 10 mM L-histidine, or GAM + 10 mM urocanate and maintained anaerobically at 37°C with shaking. Samples collected at 8-12 hour intervals over 70 hours were serially diluted in anaerobic PBS and plated on GAM agar. Plates were incubated at 37°C for 48–72 hours before CFU counting.

#### Identification of strains with reductive histidine pathway

To characterize reductive pathway activity and ImP production capacity, *O. symbiosa, B. coccoides, E. lenta, A. equolifaciens, P. excrementihominis*, and *F. ulcerans* strains were selected based on genomic identification of *hutH* and/or *urdA* genes. Strains were cultured anaerobically in 1 mL volumes supplemented with 500 µM L-histidine-^13^C_6_ hydrochloride monohydrate (Sigma) and 500 µM urocanate, or with media alone as a negative control. Cultures were performed in triplicate and sampled at designated intervals for LC-MS/MS quantification of histidine-^13^C_6_ consumption, urocanate depletion, ImP, and ^13^C_6_-ImP formation.

#### Identification of strains with oxidative histidine pathway

To characterize oxidative pathway activity and downstream metabolite production, strains predicted to encode the oxidative pathway (*A. intestini, B. coccoides, B. celeris, F. nucleatum, F. ulcerans, O. symbiosa*) were selected based on genomic identification of *hutH* and *hutU* genes, with methylaspartate or hydroxyglutarate pathway genes. Strains were cultured anaerobically in the presence of 1000 µM histidine-^13^C_6_. Cultures were performed in triplicate and sampled at designated intervals for LC-MS/MS quantification of histidine-^13^C_6_ consumption, ^13^C_2_-acetate production, and ^13^C_4_-butyrate production.

#### Glutamate-mediated suppression of oxidative histidine pathway activity

To characterize the kinetics of glutamate-mediated pathway suppression, overnight *O. symbiosa* cultures were diluted to OD_600_ = 0.05 in fresh medium and incubated anaerobically with either 10 mM histidine-^13^C_6_ alone or 10 mM histidine-^13^C_6_ + 10 mM L-glutamate. Cultures were sampled at 2.5, 5, 7.5, and 24 hours for measurement of substrate depletion and bacterial growth. Supernatants were analyzed by LC-MS to quantify histidine and glutamate consumption, while cell density was determined by serial dilution plating for CFU enumeration on appropriate agar.

To evaluate whether glutamate-mediated suppression of oxidative histidine metabolism was conserved across taxa, candidate strains (*A. intestini, B. coccoides, B. celeris, F. nucleatum, F. ulcerans*) were cultured in triplicate under three conditions: (1) media alone, (2) 1 mM histidine-^13^C_6_, or (3) 1 mM histidine-^13^C_6_ + 5 mM L-glutamate. Cultures were sampled at designated intervals for LC-MS/MS quantification of glutamate and histidine depletion, and ImP accumulation.

#### Co-culture competition assays

To assess between-strain histidine pathway competition, *P. excrementihominis* was prepared by overnight anaerobic growth in 30 mL flasks, followed by pelleting, washing, and resuspension to 0.1 OD_600_. Oxidative pathway strains (*O. symbiosa, Blautia coccoides, A. intestini*) were cultured separately to mid-exponential phase (6-8 hours), then harvested, washed, and standardized to 0.1 OD_600_.

Mixed cultures were initiated by combining equal volumes of *P. excrementihominis* with each partner strain in medium containing 1 mM histidine-^13^C_6_ with or without 5 mM glutamate supplementation. Cultures were maintained anaerobically and sampled at intervals for LC-MS/MS-based isotope tracing of histidine-derived metabolites.

### Transcriptomics

For transcriptomic analysis, nine overnight cultures of *O. symbiosa* were established in 50% BHI supplemented with either 10 mM histidine or 10 mM histidine plus 5 mM glutamate. Cultures were then diluted 1:100 into fresh medium of the same composition and adjusted to a starting OD_600_ of 0.01. Cells were grown anaerobically at 37°C to mid-log phase (approximately 6 h), harvested by centrifugation at 4,000 × g for 10 min, flash-frozen in a dry ice/ethanol bath, and stored at -80°C until RNA extraction. Cell pellets were thawed and total RNA from biological replicates were extracted using the Maxwell RSC instrument (Promega).

Extracted RNA was quantified using Qubit, and integrity was measured using TapeStation (Agilent Technologies). Libraries from ribo-depleted samples were constructed using the NEB’s Ultra Directional RNA library prep kit for Illumina. First, up to 500 ng total RNA was subjected to ribosomal RNA depletion (for bacteria) using NEBNext rRNA depletion kit. Ribosomal RNA-depleted samples were fragmented based on RNA integrity number (RIN). Post cDNA synthesis, Illumina compatible adapters were ligated onto the inserts and the final libraries were quality assessed using TapeStation (Agilent technologies). Libraries were normalized using library size and final library concentration (as determined by Qubit). Library concentration (ng/μL) was converted to nM to calculate dsDNA library concentration. Equimolar libraries were then pooled together at identical volumes to ensure even read distribution across all samples. Normalized libraries were then sequenced on Illumina’s NextSeq 1000/2000 at 2x100bp read length.

High-quality reads were mapped to the circularized hybrid assembled genome of HCS.1 (NCBI: CP170704), using Bowtie2 (v.2.4.5), and sorted with Samtools (v1.6). Read counts were generated using featureCounts (v2.0.1) with Bakta annotations.^52^ Gene expression was quantified as the total number of reads uniquely aligning to the reference genome, binned by annotated gene coordinates. Differential gene expression and quality control analyses were performed using DESeq2 in R with Benjamini– Hochberg false discovery rate adjustment applied for multiple testing corrections.^53^

### Fecal community experiments

#### Fecal sample collection

Human fecal samples from healthy, antibiotic-free individuals were collected under an IRB-approved protocol (University of Chicago Protocol IRB20-1384). All samples were de-identified prior to use in this study in accordance with the approved protocol. Samples were frozen in anaerobic 20% glycerol and stored at -80 °C until future use in enrichment and *ex vivo* experiments.

#### Fecal community enrichment cultures

Enrichment experiments were performed essentially as previously described.^54,55^ For each sample, 50 mM histidine or urocanate was added to a basal growth medium, containing 19.2 g/L Na_2_HPO_4_·7H_2_O, 4.5 g/L KH_2_PO_4_, 0.75 g/L NaCl, 1.5 g/L NH_4_Cl, 2 g/L sodium acetate, 1.5 g/L sodium formate, 0.1 g/L tryptone, 0.1 g/L Bacto yeast extract, 1 g/L MgSO_4_, and a 1x stock of minerals and vitamins and was adjusted to pH 6.5. The 1x minerals and vitamins stock solution contained 0.1 mg/L FeCl_2_·4H_2_O, 0.846 mg/L MnSO_4_·H_2_O, 0.028 mg/L ZnSO_4_·7H_2_O, 0.148 mg/L CaCl_2_·2H_2_O, 0.002 mg/L CuSO_4_·5H_2_O, 0.002 mg/L CoCl_2_·7H_2_O, 0.002 mg/L H_3_BO_3_, 0.002 mg/L Na_2_MoO_4_·2H_2_O, 50 mg/L NaCl, 1.2 mg/L trisodium citrate, 0.05 mg/L biotin, 0.1 mg/L D-pantothenic acid, 0.05 mg/L lipoic acid, 0.1 mg/L niacinamide, 0.1 mg/L para-aminobenzoic acid, 0.1 mg/L pyridoxal HCl, 0.05 mg/L riboflavin, 0.1 mg/L thiamine HCl, and 0.01 mg/L vitamin B12.

Homogenized fecal samples were pelleted and washed 3x in phosphate buffer saline (PBS), then resuspended in 1 mL PBS. 20 µL cell suspension was added to each enrichment culture condition and incubated for 72 hours. After 72 hours, 20 µL of each culture was used to inoculate fresh media supplemented with its respective compound. Cultures were passaged a total of 4 times. After the final passage, a portion of each condition was preserved in 20% glycerol and frozen at -80°C. The remaining culture was pelleted and processed for 16S rRNA sequencing.

#### Ex vivo fecal suspensions

Fresh mouse fecal pellets were collected during the early morning (8-9 AM) and immediately transferred into an anaerobic chamber. Fecal material was homogenized in pre-reduced brain heart infusion medium (BHIM) under anaerobic conditions to generate fecal slurries.

Human fecal samples were obtained from glycerol stocks previously collected and stored at -80°C. Frozen aliquots were thawed anaerobically, and glycerol was removed by centrifugation followed by two washes with pre-reduced PBS. Washed microbial pellets were resuspended in fresh pre-reduced BHIM to achieve comparable cell densities to mouse fecal slurries.

Fecal slurries (mouse) were aliquoted into culture tubes and supplemented with 1 mM histidine-^13^C_6_ alone or with 1 mM histidine-^13^C_6_ + 5 mM L-glutamate. For human stool, fecal slurries (humans) were aliquoted into culture tubes and supplemented with 2.5 mM histidine-^13^C_6_ alone or with 2.5 mM histidine-^13^C_6_ + 10 mM L-glutamate. All conditions were performed in triplicate. Cultures were incubated anaerobically at 37°C for 8 hours with periodic mixing. At the endpoint, samples were immediately quenched and processed for LC-MS/MS quantification of histidine-^13^C_6_ consumption, glutamate depletion, and ^13^C_6_-ImP formation.

### 16S rRNA amplicon sequencing

Genomic DNA from enrichment and fecal microbial communities was extracted using the QIAamp PowerFecal Pro DNA kit (Qiagen). Briefly, samples were suspended in a bead tube (Qiagen) along with lysis buffer and loaded on a bead mill homogenizer (Fisherbrand). Samples were then centrifuged, and the supernatant was resuspended in a reagent that effectively removed inhibitors. DNA was then purified routinely using a spin column filter membrane and quantified using Qubit. The 16S rRNA variable V4-V5 region was amplified using universal bacterial primers, 564F and 926R. Amplicons were purified using magnetic beads, then quantified and pooled at equimolar concentrations. The Qiagen QIAseq one-step amplicon library kit was used to ligate Illumina sequencing-compatible adaptors onto amplicons. Reads were sequenced on an Illumina MiSeq platform to generate 2 x 250 base pair reads, with 5,000-10,000 reads per sample. Amplified 16S rRNA amplicons were processed using the DADA2 pipeline in R. Forward reads were trimmed at 210 bp and reverse reads at 150 bp, to remove low-quality nucleotides. Chimeras were detected and removed using default parameters. Amplicon sequence variants between 300 and 360 bp in length were taxonomically assigned to the genus level using the RDP Classifier (v2.13) with a minimum bootstrap confidence score of 80.

### Histidine pathway abundance across disease states

Non-infant samples from daSilva *et al*. were filtered by health state (Healthy = 5852, Obesity= 829, Type 2 diabetes = 232).^56^ Taxa were grouped based on the presence of relevant metabolic pathways as determined above (species representatives were updated to UHGG (v.2.01) genomes). The relative abundance and number of species were then calculated for each sample within each group. Statistical differences between groups were calculated using the non-parametric Wilcoxon Rank Sums Test and corrected for multiple testing using the Benjamini-Hochberg method.

### Mouse studies

#### Mouse husbandry

All procedures were approved by the University of Chicago Institutional Animal Care and Use Committee and performed in accordance with institutional ethical guidelines. Germ-free C57BL/6J mice were maintained in sterile flexible film isolators at the Gnotobiotic Research Animal Facility. SPF male C57BL/6J mice were acquired from The Jackson Laboratory at 4 weeks of age and housed under barrier conditions with 12-hour light/dark cycles and unrestricted access to chow and water.

#### Mouse metabolic phenotyping

For glucose tolerance testing, mice underwent a 6-hour fast before receiving intraperitoneal glucose (1 or 2 g/kg body weight). Tail vein blood glucose was monitored at baseline and at 10, 20, 30, 60, 90, and 120 minutes post-injection using a handheld glucometer (AlphaTRAK 3). Area under the curve (AUC) was calculated using GraphPad Prism version 11.0.0 for statistical comparison between treatment groups.

#### Mouse gnotobiotic monocolonization model

Germ-free male C57BL/6J mice (8-12 weeks old) received a single oral gavage of *Fusobacterium ulcerans, Parasutterella excrementihominis*, or *Escherichia coli* and were maintained under gnotobiotic conditions throughout the experiment. Following a 7-day colonization period, mice were fasted for 6 hours beginning at light onset. Blood was collected via submandibular bleeding at the start of the fasting period. Following the fast, mice underwent intraperitoneal glucose tolerance testing. Animals were euthanized and cecal contents were collected for metabolomic profiling and microbial community analysis. Colonization was verified through culture-based enumeration and/or 16S rRNA gene sequencing.

#### HFD-consortia gnotobiotic mouse model

Germ-free male C57BL/6J mice (8-12 weeks old) were transitioned to high-fat diet (Teklad Custom Diet, TD.96132) And maintained under gnotobiotic conditions to establish diet-induced metabolic dysfunction. After 8 weeks of HFD feeding, mice were colonized by oral gavage with a previously characterized “base consortia” of 12 human gut bacterial species lacking histidine metabolic pathways (**Table S7**).^35^

To evaluate the impact of histidine pathway activity on ImP production and glucose metabolism, experimental groups received: (1) base consortia alone (control), (2) base consortia + *F. ulcerans* (reductive pathway), (3) base consortia + *O. symbiosa* (oxidative pathway), or (4) base consortia + *F. ulcerans* + *O. symbiosa* (pathway competition). Colonization was permitted to stabilize for 14 days before mice underwent fasting blood collection followed immediately by glucose tolerance testing as described above. Following test completion, mice were euthanized and cecal contents were harvested for metabolomic profiling and 16S rRNA amplicon sequencing to confirm community composition.

#### MSG supplementation and microbiome depletion in SPF mice

Specific pathogen-free male C57BL/6J mice (4 weeks old) were obtained from The Jackson Laboratory and transitioned to high-fat diet (Research Diets, D12492i) under barrier conditions. After 6 weeks of HFD feeding to establish metabolic dysfunction, mice were assigned to antibiotic-treated or untreated groups.

For microbiome depletion, mice received sterile drinking water containing a broad-spectrum antibiotic cocktail: ampicillin (1 g/L), neomycin sulfate (1 g/L), metronidazole (1 g/L), and vancomycin (0.5 g/L). Solutions were prepared aseptically with metronidazole dissolved by brief heating and vortexing. Antibiotic water was shielded from light and replaced every 48-72 hours to maintain efficacy. Treatment commenced 7 days prior to dietary intervention and continued throughout the experimental period. Fecal samples were collected at baseline and during treatment to verify microbial depletion by culture-based enumeration.

Following the 7-day antibiotic pretreatment (or equivalent timeline for untreated controls), mice were maintained on HFD and provided drinking water supplemented with either 1% (w/w) MSG (59 mM) + 0.0155% (w/w) histidine (1 mM), or 0.0155% (w/w) histidine (1 mM) + equimolar NaCl (59 mM) as a sodium-matched control. MSG dose was based on concentrations used in prior metabolic studies.^39^

After 7 days of MSG or NaCl supplementation, mice were fasted for 6 hours beginning at light onset. Blood was collected via submandibular bleeding at the start of the fasting period. Following the fast, mice underwent intraperitoneal glucose tolerance testing. Upon completion of the assay, animals were euthanized and cecal contents were collected for metabolomic and microbial analyses. Body weight, food intake, and water consumption were monitored throughout the study.

#### Dietary MSG crossover intervention in SPF mice

Specific pathogen-free male C57BL/6J mice (4 weeks old) were obtained from The Jackson Laboratory and transitioned to high-fat diet (Research Diets, D12492i) under barrier conditions. Following 6 weeks of HFD feeding to induce metabolic dysfunction, mice were randomly allocated to receive drinking water supplemented with either 1% (w/v) MSG (59 mM) + 0.0155% (w/v) histidine (1 mM), or 0.0155% (w/v) histidine (1 mM) + equimolar NaCl (59 mM) as a sodium-matched control, while continuing on HFD. After 7 days of MSG or NaCl supplementation, mice were fasted for 6 h beginning at light onset. Blood was collected via submandibular bleeding at the start of the fasting period. Following the fast, mice underwent intraperitoneal glucose tolerance testing. Mice were then returned to unsupplemented HFD for a 7-day washout period before being switched to the reciprocal dietary condition (MSG→control or control→MSG) for an additional 7 days. Following this second treatment period, fasting blood collection and glucose tolerance assessment were repeated. Upon completion of the final glucose tolerance test, mice were euthanized and cecal contents collected for metabolomic profiling and microbial community analysis.

### Metabolite extraction and quantification

#### Cecal metabolite extraction

Cecal contents were weighed and transferred to bead-beating tubes. Samples were homogenized to generate a working homogenate at a final concentration of 100 mg/mL. For most samples, 500 µL of working homogenate was prepared, corresponding to 50 mg-equivalent of cecal material. For smaller samples, 250 µL of working homogenate was prepared (25 mg-equivalent).

For metabolite extraction, 10 µL of the 100 mg/mL homogenate (1 mg-equivalent) was diluted with 90 µL water. Four hundred microliters of cold acetonitrile containing internal standards was added, and samples were vortexed and centrifuged to precipitate proteins. Four hundred microliters of supernatant was transferred to a new tube and dried under vacuum. Dried extracts were derivatized by butylation, dried again, and resuspended in 80 µL solvent for LC-MS/MS analysis.

#### Serum metabolite extraction

Serum samples were diluted 1:5 in water prior to extraction. Following dilution, 400 µL of cold acetonitrile containing internal standards was added. Samples were vortexed and centrifuged, and supernatants were transferred and dried under vacuum. Dried extracts were subjected to butylation derivatization, dried again, and resuspended in 80 µL solvent prior to LC–MS/MS analysis.

#### LC-MS/MS quantification of ImP, amino acids, and urocanate

LC-MS quantification was performed on an Agilent 6460 Triple Quadrupole LC/MS system coupled to a 1290 UHPLC (Agilent Technologies), operated in positive electrospray ionization (ESI+) mode. Compounds were separated on an ACQUITY UPLC HSS T3 column (2.1 × 100 mm, 1.8 µm; Waters) using a linear gradient (flow rate: 0.4 mL/min) of (A) water with 0.1% formic acid and (B) acetonitrile with 0.1% formic acid, ramping from 0% to 60% B over 4 min at 40 °C. The following transitions were monitored in dynamic MRM mode: glutamate-d_5_ (*m/z* 265 → 135), glutamate (*m/z* 260 → 129.9), [U-^13^C]-histidine (*m/z* 218 → 115), histidine (*m/z* 212 → 110), cinnamate (*m/z* 205 → 131), [U-^13^C]-imidazole propionate (ImP) (*m/z* 203 → 128.7), imidazole propionate (ImP) (*m/z* 197 → 123.1), and urocanate (*m/z* 195 → 121), all detected as [M+H]^+^ ions. Instrument settings were as follows: capillary voltage 3500 V, nozzle voltage 500 V, gas temperature 300 °C, and gas flow 10 L/min.

#### Isotope-labeled SCFA and glutamate quantification by LC-MS

For the experiment measuring incorporation of isotopically labeled carbons from histidine-^13^C_6_ into glutamate, butyrate, and acetate in *O. symbiosa* and *B*.*coccoides* cultures, extractions were performed using LC-MS grade methanol (Fisher Scientific; Optima A456) with 500 µM sodium D_7_-butyrate (98%, Cambridge Isotope Laboratories; DLM-7616) and sodium D_3_-acetate (Cambridge Isotope Laboratories, DLM-3126-25) as internal standards (ISTDs). An L-glutamic acid standard (Sigma-Aldrich; G8415) was analyzed separately to confirm glutamate ions. Derivatization reagents were *N*-(3-dimethylaminopropyl)-*N*′-ethylcarbodiimide hydrochloride (EDC; Sigma-Aldrich; E7750) and 3-nitrophenylhydrazine hydrochloride (3-NPH; Sigma-Aldrich; N21804). Solvents used were LC-MS grade water (Fisher Scientific; Optima W7), LC-MS grade acetonitrile (Fisher Scientific; Optima A955), and 99% Ultra-Pure LC-MS Grade formic acid (CovaChem; 11202) as a mobile phase buffer.

Culture supernatants were stored at -80 °C prior to LC-MS analysis. Four volumes of methanol containing ISTDs were added to one volume of each culture supernatant in a microcentrifuge tube, followed by vortexing for 30 seconds. The tubes were then centrifuged at −10 °C and 20,000 × *g* for 15 min. Based on methods from Mullowney, *et al*., 25 µL of extract supernatant from each sample was transferred to a mass spectrometry vial with 12.5 µL (200 mM) 3-nitrophenylhydrazine (3-NPH) and 12.5 µL (120 mM) N-(3-dimethylaminopropyl)-N′-ethylcarbodiimide hydrochloride.^57^ Samples were incubated at 40 °C for 30 minutes and then chilled at -80 °C for two minutes. LC–MS analysis was performed using a Thermo Scientific Vanquish Flex UHPLC coupled to an IQ-X mass spectrometer (Thermo Fisher Scientific). Sample injections of 3 µL were separated on a CORTECS T3 column (Waters, 120Å, 1.6 µm, 2.1 x 100 mm) fitted with a KrudKatcher Ultra Column In-Line Filter (Phenomenex, AF0-8497), and a CORTECS T3 VanGuard pre-column (Waters, 120Å, 1.6 µm, 2.1 mm X 5 mm) at 40°C with a flow rate of 0.350 mL/min and run-time of 15 min. The aqueous mobile phase was water and the organic mobile phase was acetonitrile, both supplemented with 0.1% formic acid. The chromatographic method used was isocratic 5% acetonitrile for 1.0 min, then a gradient of 5 to 100% acetonitrile for 10.0 min, a wash of 100% acetonitrile for 3 min, a 0.1 min decrease from 100 to 5% acetonitrile, and a 1.9 min re-equilibration at 5% acetonitrile. Flow from the UHPLC was negatively ionized with the HESI source set to 3,500 V, ion transfer tube temperature set to 190 °C, vaporizer temperature set to 120 °C, and sheath, auxiliary and sweep gases set to arbitrary values of 50, 10 and 1, respectively. Centroid data for MS^1^ were acquired using a maximum injection time of 50 ms, 1 microscan, a radio frequency (RF) lens value of 35 %, a normalized automatic gain control (AGC) target of 25%, an *m/z* 133–2,000 quadrupole isolation scan range and an Orbitrap resolution of 120,000.

Glutamic acid readily cyclizes to form pyroglutamic acid both under the EDC-catalyzed 3-NPH derivatization conditions and during electrospray ionization in LC-MS analysis.^58^ Regardless of the isotopic labeling, no singly derivatized glutamic acid ions were observed and doubly derivatized glutamic acid levels were low in abundance in the glutamic acid standard and experimental samples. Thus, ^13^C_5_-glutamic acid in experimental samples was measured as the highly abundant 3-NPH-^13^C_5_-pyroglutamic acid, while 3-NPH-pyroglutamic acid derived from the glutamic acid standard was used for retention time comparison. 3-NPH-^13^C_4_-butyrate and 3-NPH-^13^C_2_-acetate retention times were confirmed by comparison to 3-NPH-d_7_-butyrate and 3-NPH-d_3_-acetate, respectively, both arising from the ISTDs in the methanol extraction.

## ACKNOWLEDGEMENTS

Research reported in this publication was supported by funding from the Arnold and Mabel Beckman Foundation (Beckman Young Investigator to M.M.), the Brinson Foundation (to S.H.L), the Deutsche Forschungsgemeinschaft German Research Foundation (542537779 to C.J.), and the National Institutes of Health (T32 GM150375 to T.H.B, T32 GM144292 to I.T.Y., R01 DK060581 to R.G.M, P30 DK020595 to R.G.M, R35 GM147478 to M.M., and R35 GM146969 to S.H.L). We thank Shanna Banogon, the Duchossois Family Institute, and University of Chicago Diabetes Research and Training Center for funding and experimental support.

## SUPPLEMENTAL MATERIAL

## Supplemental Tables

Table S1. *O. symbiosa* normalized transcript counts when grown on histidine.

Table S2. Histidine pathway predictions for UHGG species representatives.

Table S3. Association between culture histidine depletion and oxidative histidine pathway reported by Han et al.

Table S4. Taxonomic composition and histidine pathway assignments for metagenomes in Figures S6 and S7.

Table S5. Differential gene expression in *O. symbiosa* grown on histidine with or without glutamate.

Table S6. Reference enzymes and homology thresholds for histidine pathway identification.

Table S7. Strains used in this study

**Figure S1.**
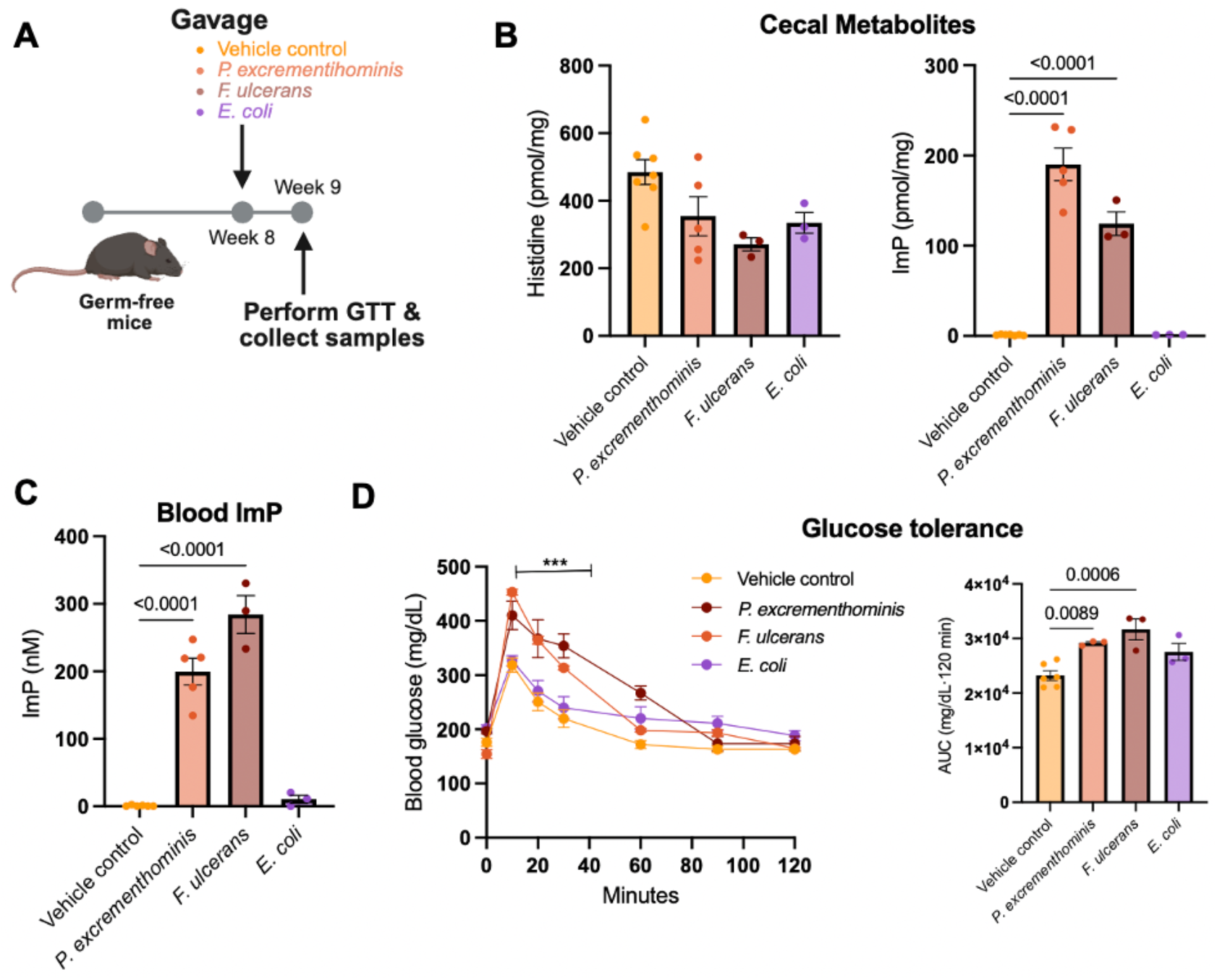
Monocolonization with reductive histidine pathway bacteria elevates systemic ImP and impairs glucose tolerance. (A) Experimental design for gnotobiotic monocolonization study. Mice were administered *F. ulcerans, P. excrementihominis, E. coli* (histidine pathway-lacking control), or vehicle (PBS). (B) Cecal metabolite concentrations determined by targeted LC-MS. (C) Blood ImP concentration. (D) Glucose tolerance assessed by intraperitoneal glucose tolerance test. All data shown as mean ± SEM and analyzed by one-way ANOVA with Tukey’s post-hoc test.

**Figure S2.**
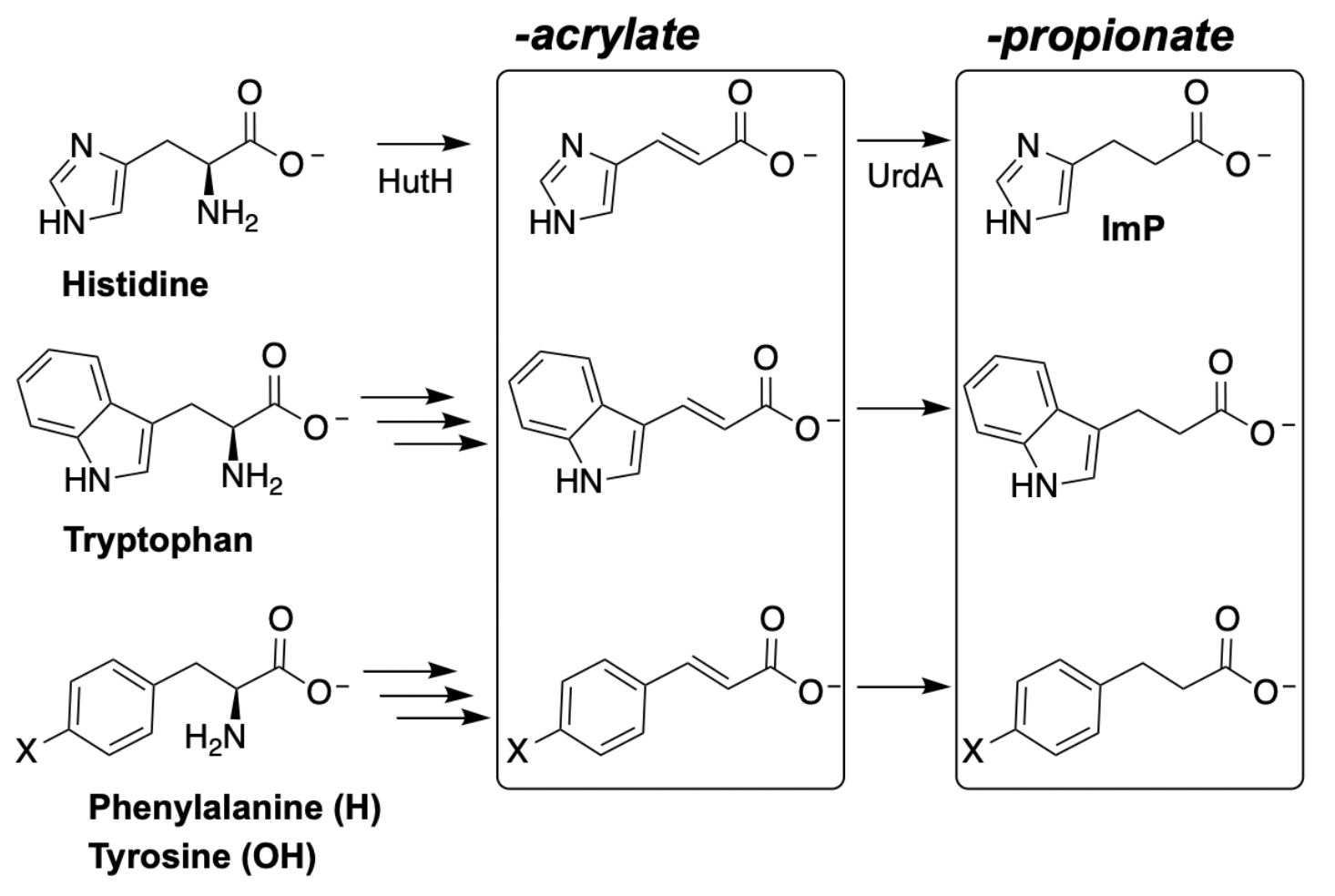
The reductive histidine pathway shares structural and mechanistic features with characterized reductive Stickland pathways. Comparison of histidine metabolism via HutH and UrdA with characterized phenylalanine, tyrosine, and tryptophan Stickland pathways. Conserved features include: (1) amino acid deamination to α,β-unsaturated acrylate intermediates, (2) reductase-mediated reduction to saturated products, and (3) generation of “-propionate” carboxylate products.

**Figure S3.**
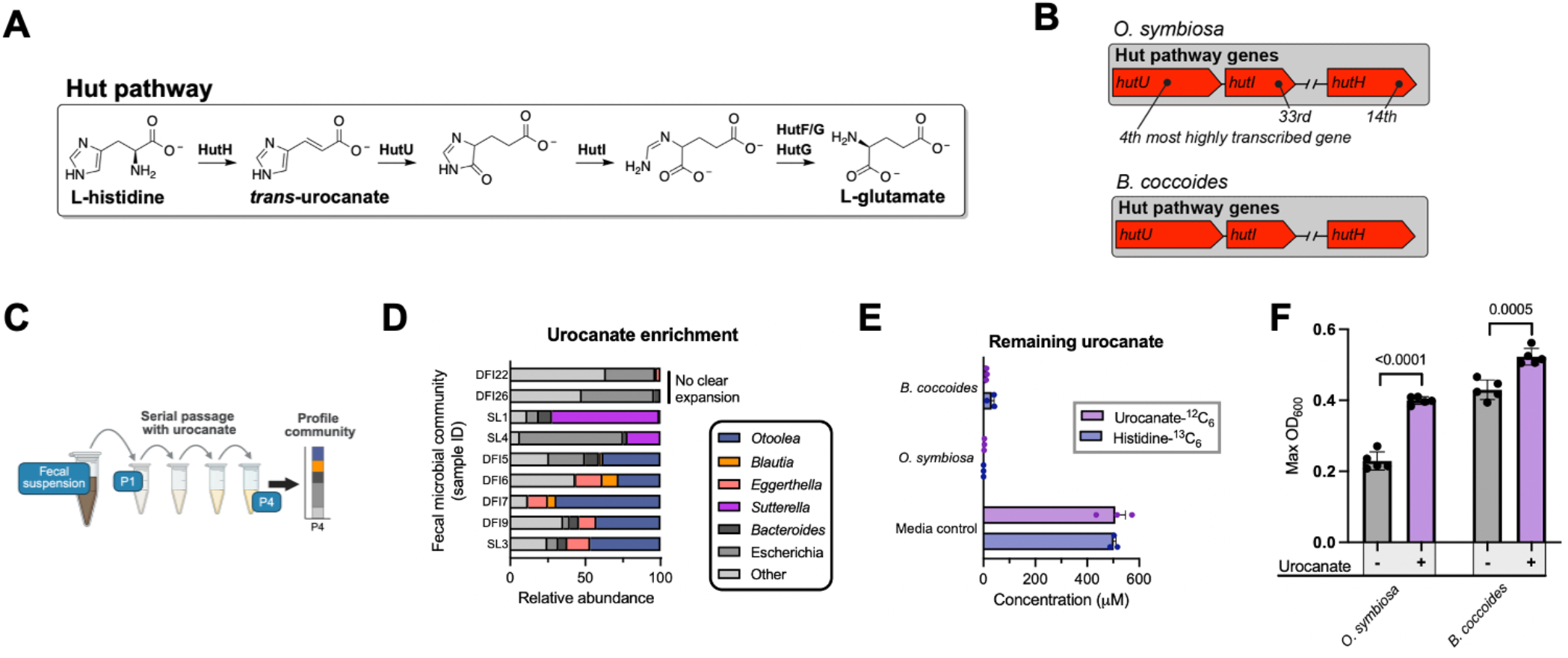
Evidence for Hut pathway-mediated histidine utilization in *O. symbiosa* and *B. coccoides*. (A) Enzymatic steps in histidine to glutamate conversion via the Hut pathway. The enzymes that catalyze the final step (HutG and/or HutF) in the pathway exhibit considerable species-level variability and consequently were not analyzed in our studies. (B) Genomic organization of Hut pathway genes in *O. symbiosa* and *B. coccoides*. Gene expression rankings based on normalized transcript abundance from 4,175 predicted *O. symbiosa* open reading frames from RNA-seq analysis of *O. symbiosa* cultured with histidine as the primary carbon source. (C) Experimental design for enriching urocanate-catabolizing bacteria from human fecal communities through serial passaging with urocanate as the primary substrate. (D) Community composition of human fecal microbial communities following urocanate enrichment passaging. (E) Remaining unlabeled urocanate and histidine-^13^C_6_ in culture media of *O. symbiosa* and *B. coccoides* supplemented with both substrates. (F) Growth of *O. symbiosa* and *B. coccoides* cultured with or without 10 mM urocanate supplementation. Data in E-F shown as mean ± SEM and analyzed by Two-tailed Welch’s t-test.

**Figure S4.**
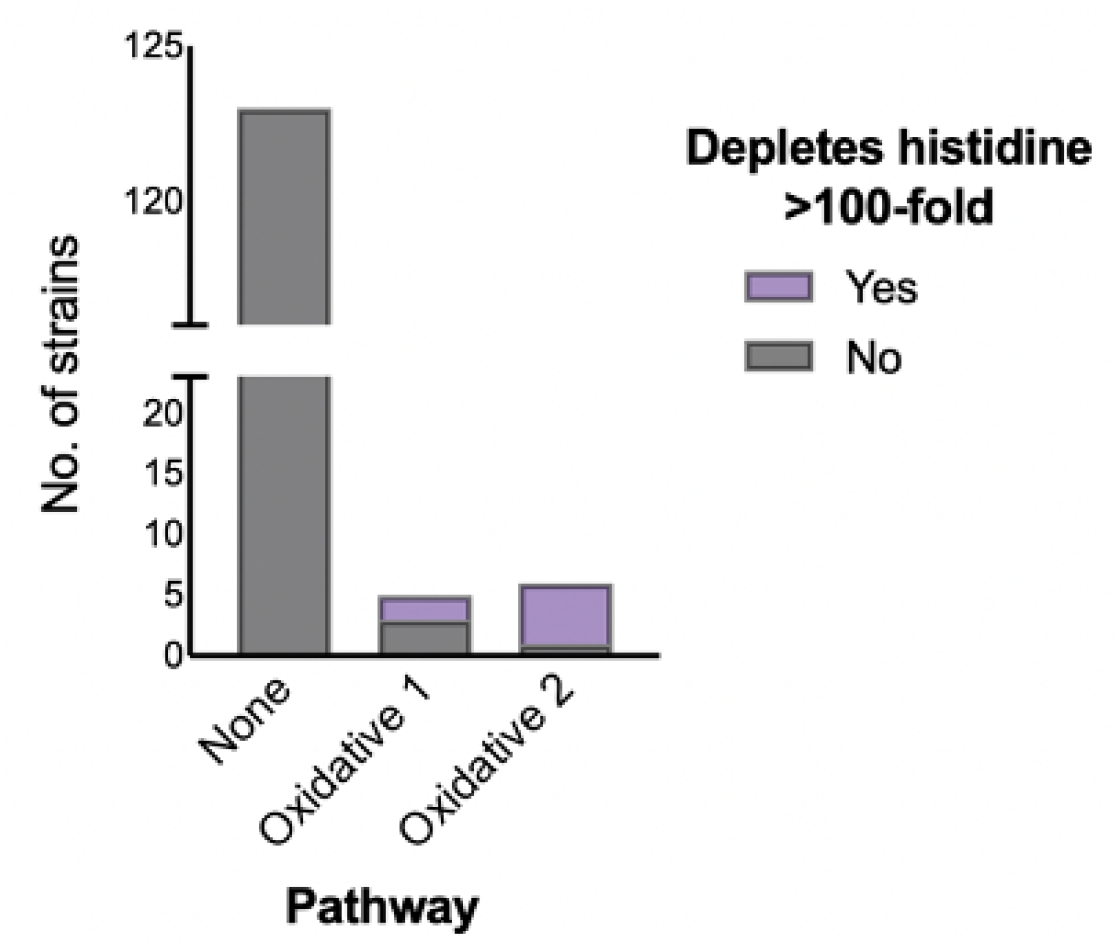
Histidine-depleting activity correlates with oxidative histidine pathway gene content in gut bacteria. Analysis of published metabolomics data (Han et al.) quantifying histidine depletion from culture media of individually grown gut bacterial strains. Strains are grouped by pathway genomic analysis of gene content: (1) none (no oxidative histidine pathway genes); (2) oxidative 1 (Hut pathway and methylaspartate pathway genes); (3) oxidative 2 (Hut pathway and 2-hydroxyglutarate pathway genes). Bars indicate the number of strains in each group that substantially depleted histidine (>100-fold reduction) from culture media. See **Table S3** for strain identities.

**Figure S5.**
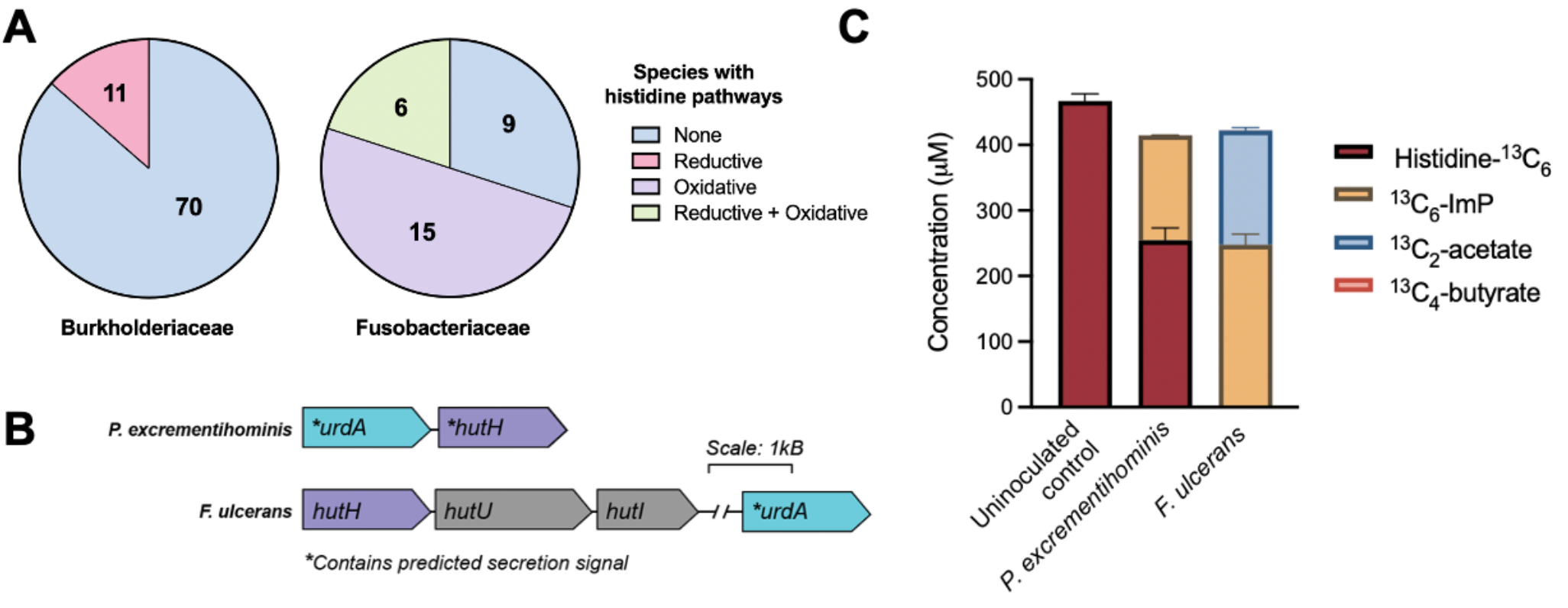
Distinct genomic organization of the reductive histidine pathway in reductive pathway-encoding bacteria. (A) Histidine pathway gene content in UHGG genomes from reductive pathway-encoding families Burkholderiaceae and Fusobacteriaceae. Candidate oxidative and reductive histidine pathways were identified using marker genes, with genomes categorized as: (1) None (no pathway genes), (2) Reductive (Reductive pathway only), (3) Oxidative (Oxidative pathway only), or (4) Reductive + Oxidative (both reductive and oxidative pathway). Genome counts per category are shown. See **Table S2** for strain identities. (B) Representative genetic loci showing reductive histidine pathway gene organization in Burkholderiaceae (*P. excrementihominis*) and Fusobacteriaceae (*F. ulcerans*). Burkholderiaceae lack the oxidative histidine pathway and *urdA* and *hutH* co-localize. In Fusobacteriaceae, *hutH* co-localizes with oxidative histidine pathway genes (*hutU* and *hutI*), while *urdA* occupies a separate genomic location. (C) ^13^C-labeled metabolite production from *P. excrementihominis* and *F. ulcerans* cultures supplemented with histidine-^13^C_6_.

**Figure S6.**
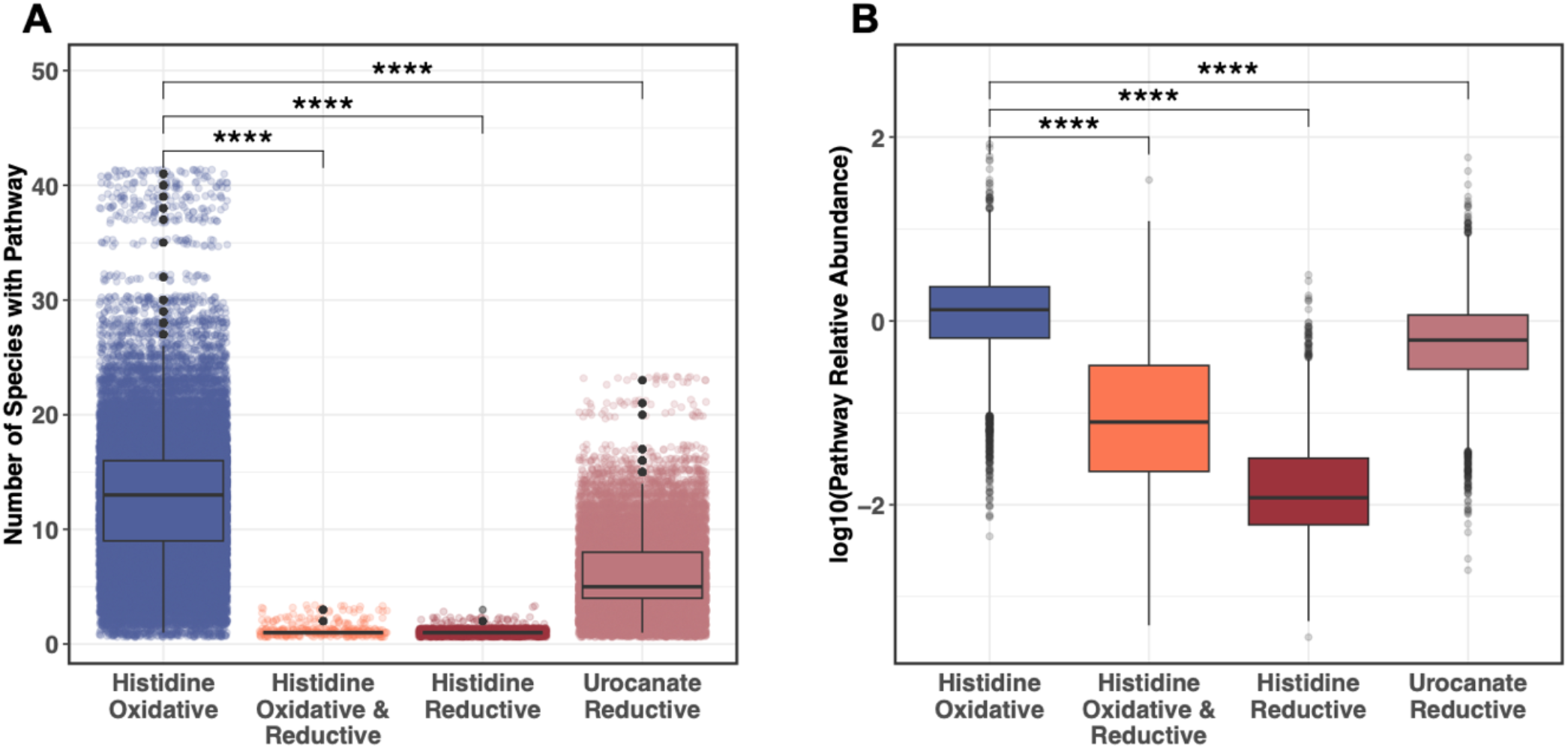
The oxidative histidine pathway is more prevalent and abundant than the reductive pathway in human gut microbiomes. Analysis of publicly available human fecal metagenomes showing distribution of bacterial histidine pathway types. Categories represent mutually exclusive pathway configurations as designated in **Table S2**: (1) Oxidative: species encoding Hut and glutamate fermentative pathways only; (2) Reductive: species encoding only reductive histidine pathway genes; (3) Oxidative & Reductive: species encoding genes for both pathways; (4) Urocanate Reductive: species encoding *urdA* without *hutH*, which potentially convert urocanate but not histidine to ImP. (A) Number of bacterial species in each pathway category per metagenome. Each point represents one metagenome. (B) Cumulative relative abundance of species in each pathway category per metagenome, calculated by summing the relative abundances of all species within each category. Statistical analysis by Wilcoxon rank-sum test with Benjamini-Hochberg correction for multiple testing.

**Figure S7.**
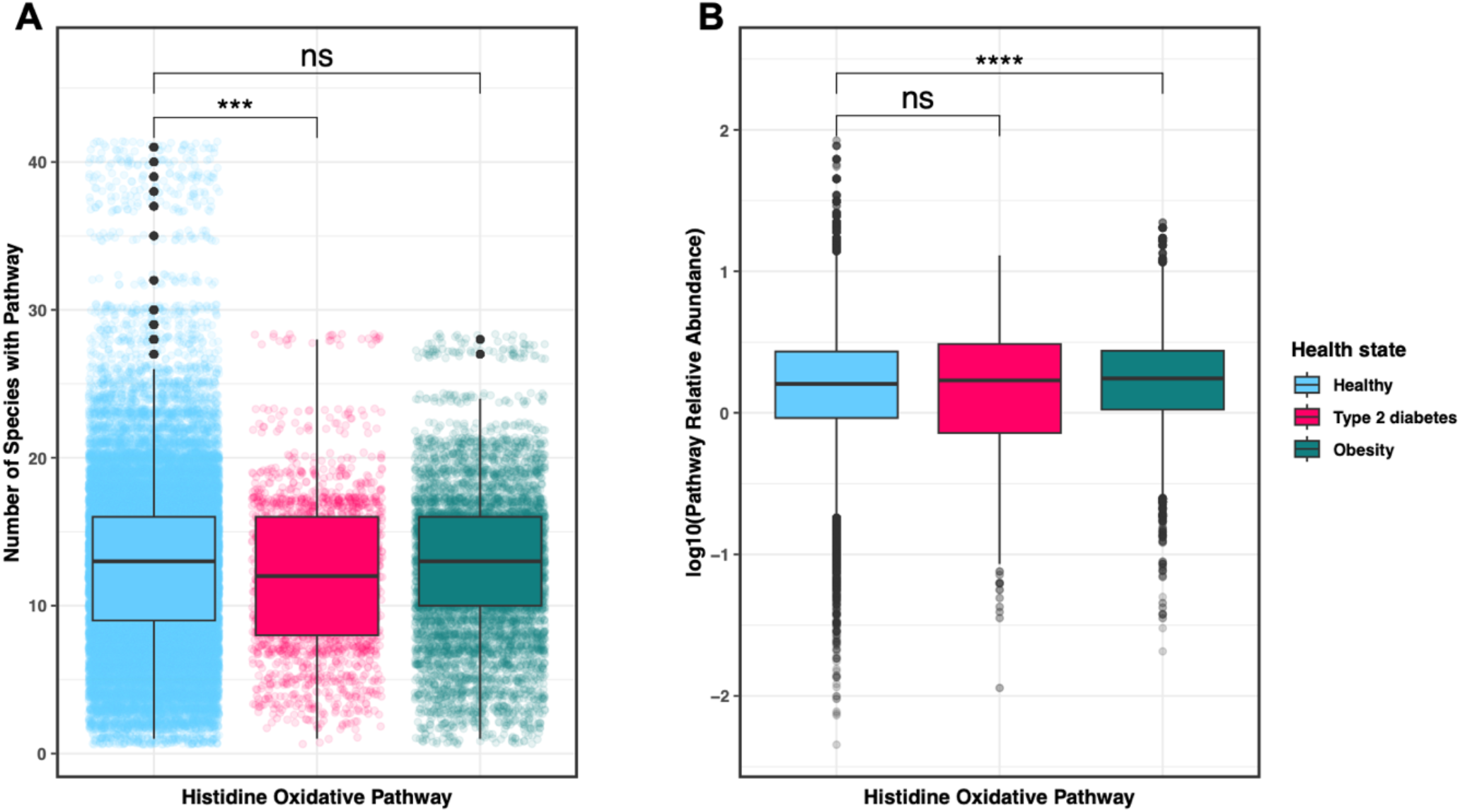
Oxidative histidine pathway bacteria show similar prevalence and abundance in healthy and cardiometabolic disease cohorts. (A) Analysis of fecal metagenomes from healthy controls, type 2 diabetes patients, and obese individuals, showing the number of bacterial species encoding oxidative histidine pathways detected per metagenome. Each point represents one individual. (B) Cumulative relative abundance of all oxidative pathway species per metagenome, calculated by summing the relative abundances of all species encoding the oxidative pathway within each sample. Statistical analysis by Wilcoxon rank-sum test with Benjamini-Hochberg correction for multiple testing.

**Figure S8.**
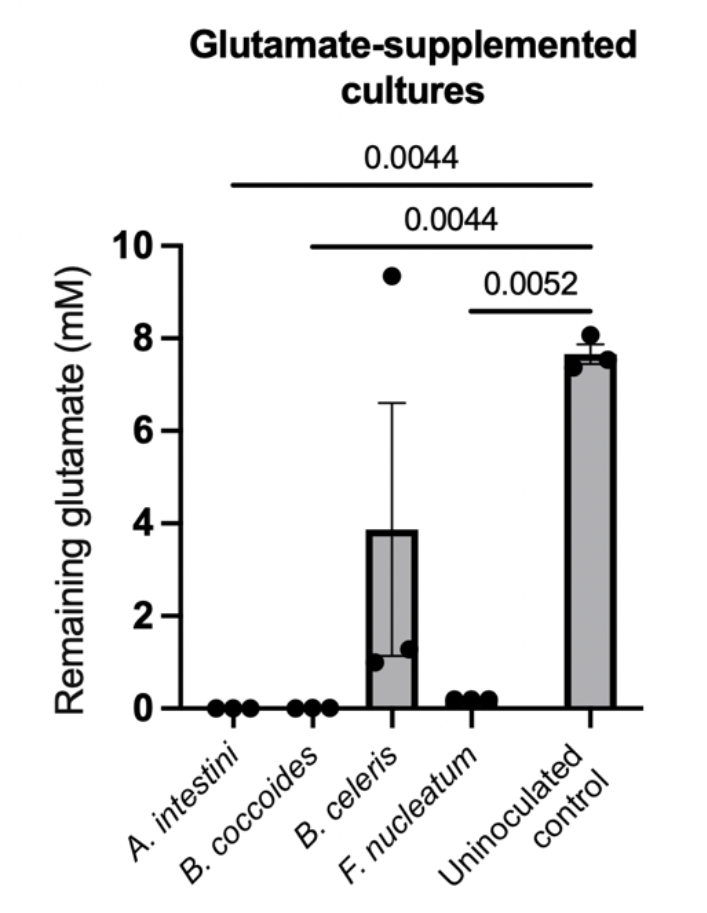
Oxidative pathway bacteria efficiently deplete glutamate while suppressing histidine consumption. Remaining glutamate in cultures of oxidative histidine pathway incubated with histidine-^13^C_6_ + glutamate, from conditions showing glutamate-mediated histidine consumption suppression shown in **Figure 3E**.

**Figure S9.**
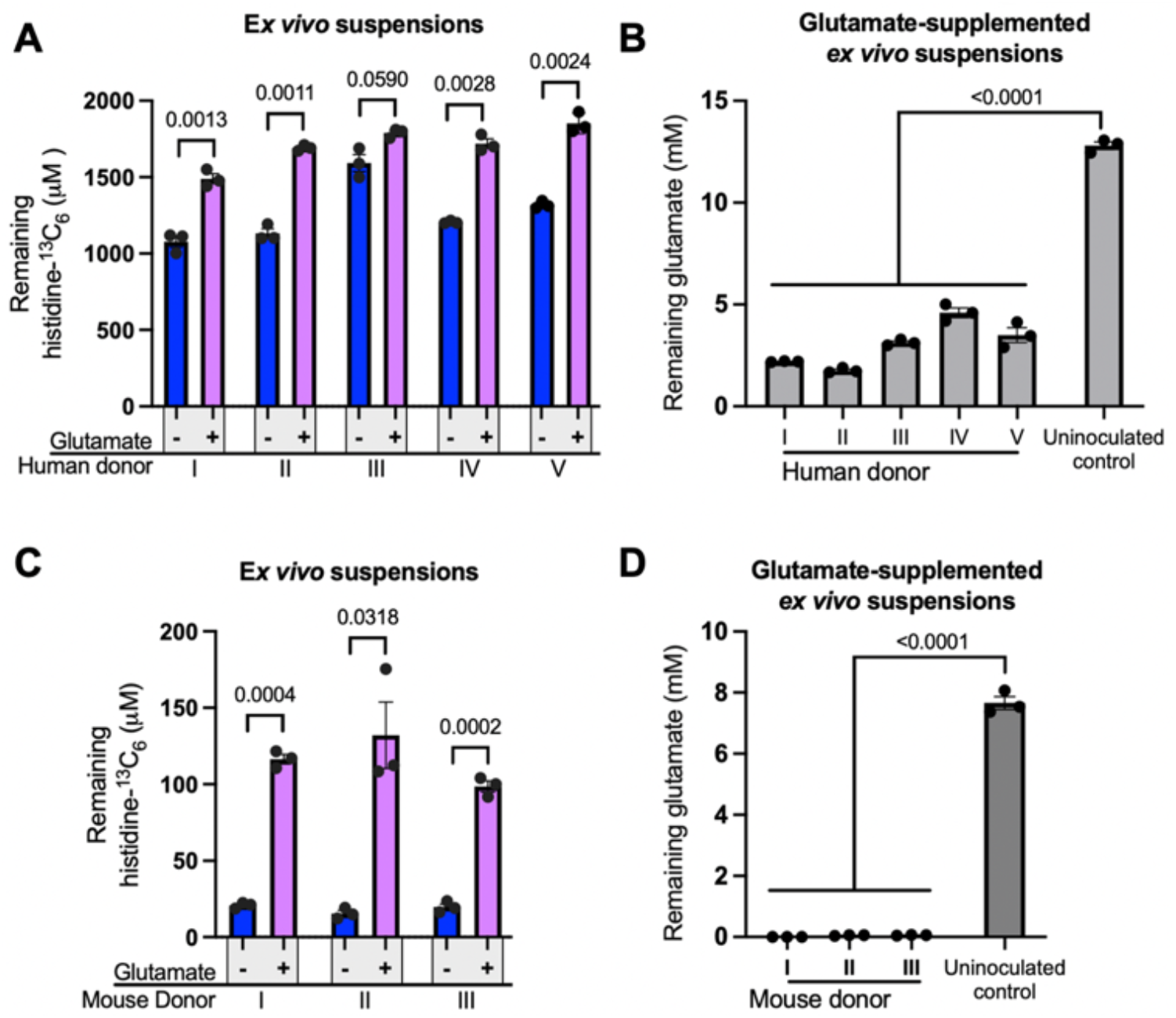
Glutamate consumption coincides with suppressed histidine depletion in natural microbial communities. Substrate consumption data from the ex vivo incubations shown in **Figure 4E** (human communities) and **Figure 4F** (mouse communities). (A) Remaining histidine-^13^C_6_ following *ex vivo* incubation of human fecal suspensions with histidine-^13^C_6_ ± glutamate. (B) Remaining glutamate following *ex vivo* incubation of human fecal suspensions with histidine-^13^C_6_ + glutamate. (C) Remaining histidine-^13^C_6_ following *ex vivo* incubation of mouse fecal suspensions with histidine-^13^C_6_ ± glutamate. (D) Remaining glutamate following *ex vivo* incubation of mouse fecal suspensions with histidine-^13^C_6_ + glutamate. All data shown as mean ± SEM and analyzed by Two-tailed Welch’s t-test or one-way ANOVA with Tukey’s post-hoc test.

**Figure S10.**
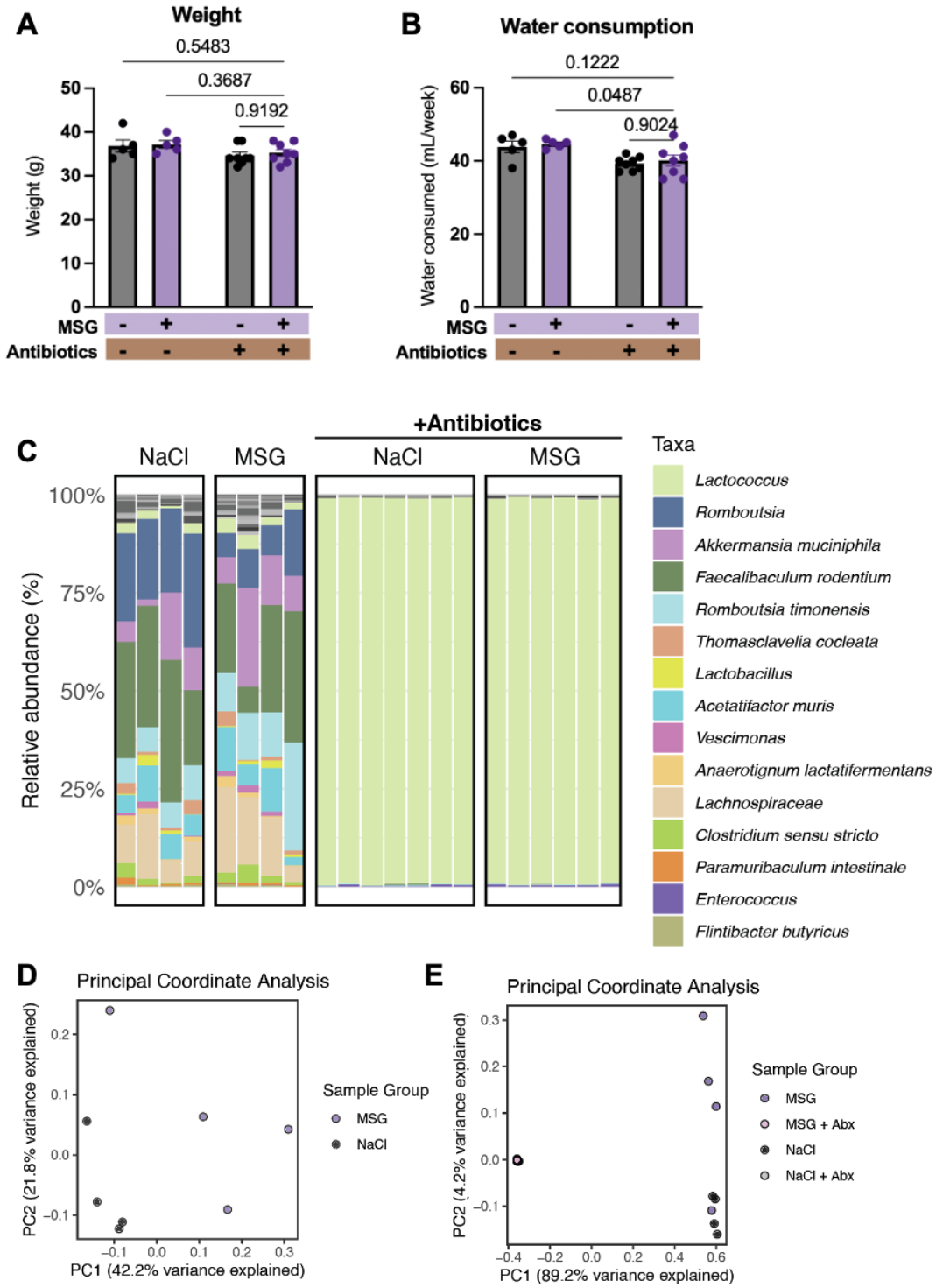
MSG supplementation has no major effects on body weight, water consumption, or microbiome composition. Physiological and microbiome data from the mouse MSG experiment shown in **Figure 5**. (A) Body weight and (B) weekly water consumption in mice receiving MSG or equimolar NaCl via drinking water with or without antibiotic treatment (Abx). Data in A-B shown as mean ± SEM and analyzed by one-way ANOVA. (C) Bacterial community composition determined by 16S rRNA amplicon sequencing. (D) Principal coordinate analysis of fecal 16S rRNA profiles based on Bray-Curtis dissimilarity of non-antibiotic-treated samples only, showing no significant separation between MSG and NaCl groups (Two-tailed Welch’s t-test, P > 0.05). (E) Principal coordinate analysis (PCoA) of fecal 16S rRNA profiles based on Bray-Curtis dissimilarity, showing all treatment groups.

